# VISTA Uncovers Missing Gene Expression and Spatial-induced Information for Spatial Transcriptomic Data Analysis

**DOI:** 10.1101/2024.08.26.609718

**Authors:** Tianyu Liu, Yingxin Lin, Xiao Luo, Yizhou Sun, Hongyu Zhao

## Abstract

Characterizing cell activities within a spatially resolved context is essential to enhance our understanding of spatially-induced cellular states and features. While single-cell RNA-seq (scRNA-seq) offers comprehensive profiling of cells within a tissue, it fails to capture spatial context. Conversely, subcellular spatial transcriptomics (SST) technologies provide high-resolution spatial profiles of gene expression, yet their utility is constrained by the limited number of genes they can simultaneously profile. To address this limitation, we introduce VISTA, a novel approach designed to predict the expression levels of unobserved genes specifically tailored for SST data. VISTA jointly models scRNA-seq data and SST data based on variational inference and geometric deep learning, and incorporates uncertainty quantification. Using four SST datasets, we demonstrate VISTA’s superior performance in imputation and in analyzing large-scale SST datasets with satisfactory time efficiency and memory consumption. The imputation of VISTA enables a multitude of downstream applications, including the detection of new spatially variable genes, the discovery of novel ligand-receptor interactions, the inference of spatial RNA velocity, the generation for spatial transcriptomics with in-silico perturbation, and an improved decomposition of spatial and intrinsic variations.

## 1 Introduction

Single-cell RNA sequencing (scRNA-seq) has matured into a robust technology that allows for the measurement of the complete gene expression profile of individual cells within various tissues [1–3]. Despite its advancements, scRNA-seq does not account for the spatial positioning of cells, a critical aspect known as spatial information. To bridge this gap, spatial transcriptomic technologies have been developed, enabling the investigation of the cellular spatial context [4]. These technologies fall into two primary categories: (1) Imaging-based methods, including smFISH [5, 6], MERFISH [7], seq-FISH [8] and Xenium [9, 10]; and (2) Sequencing-based methods , including STARmap [11] and Slide-seq [12]. Compared to spot-based spatial technologies [4, 13], these methods facilitate the capture of gene expression at a single-molecule/subcellular resolution, which can then be aggregated to deduce gene expression levels and cell structure in single-cell resolution. The analysis of single-cell gene expression enriched with spatial information holds great promise for understanding spatial-induced information.

However, the scope of genes sequenced by imaging-based and sequencing-based spatial transcriptomic methods is notably constrained, typically capturing approximately 30 to 10,000 genes [5, 7, 9, 14]. This is considerably less than the roughly 20,000 to 30,000 genes profiled by scRNA-seq [3]. To address this limitation, researchers have developed methodologies for the prediction of unmeasured gene expression within these datasets by aggregating nearest neighbors of cells for regression [15–18], joint probabilistic modeling [19–21], and transport [22–24]. These methods have been thoroughly benchmarked [10, 25]. These imputation tools utilize scRNA-seq data from the corresponding tissue to enhance the spatial transcriptomic data, and the results can then be employed in downstream analyses such as the identification of novel spatially variable (SV) genes [26] and the examination of cell-cell interactions (CCIs) [27].

Despite these progresses, there are limitations in the existing methods for imputing spatial transcriptomic data in terms of both performance and utility in practice. One primary issue is their subpar imputation quality for large-scale datasets. For instance, datasets sequenced using Xenium, which may contain more than 100,000 cells, yield a Spearman correlation coefficient below 0.5 in validation sets for mouse brain data [10]. These methods also lack scalability when applied to large-scale datasets. Tangram, for example, suffers from out-of-memory (OOM) errors when applied to large-scale datasets (smaller than 10,000 cells), and SpaGE exhibits excessively long runtime (more than 40,000 seconds). While gimVI can accommodate large-scale datasets by adjusting the batch size, it does not model spatial information and necessitates a compromise between scalability and running time as the smaller the batch size is, the longer time we need for the same device. Additionally, none of the methods sufficiently account for the discrepancy between reference scRNA-seq data and the target spatial transcriptomic data. Consequently, there is a clear need for more in-depth analysis of the existing methods to overcome the drawbacks in their designs and to develop a more effective approach to the imputation challenge.

At the same time, few methods can estimate the uncertainty of gene expression levels and discuss the effect of prediction errors on downstream applications. TransImp [28] estimates the uncertainty based on a post-imputation regression, which requires extra fitting steps. TISSUE [29] states that it can produce well-calibrated uncertainty measures for gene expression prediction, but it needs to separate samples to create the calibration set and select genes with high certainty.

To bridge the gap between imputation outcomes and actual gene expression data, we develop a pipeline tailored for the task of imputing spatial transcriptomic data. This process involves a thorough analysis of the reference scRNA-seq and target spatial data pairs, as well as the integration of spatial information during model training. Our proposed solution, named **VISTA**, leverages a novel joint probabilistic modeling approach. We combine a Graph Neural Network (GNN) [30, 31] architecture and graph sampling strategy [32] with **V**ariational **I**nference (VI) [33] for **S**patial **T**ranscriptomic **A**nalysis. VISTA is trained to predict missing gene expression levels with graph sampling for large-scale spatial data, using scRNA-seq data as a reference. By leveraging the contributions of highly correlated genes for training, VISTA utilizes probabilistic modeling for each cell and gene to estimate the uncertainty in our imputation results. Our uncertainty quantification method does not require sacrificing samples in the training dataset to generate calibration datasets while integrating this estimation approach directly into our pipeline. We demonstrate the performance of our proposed method through a series of experiments of imputation by evaluating it from both statistical and biological perspectives. Finally, our case studies of exploring Spatially Variable (SV) genes, Cell-Cell Interactions (CCIs), RNA velocity, and in-silico perturbations suggest that our imputed spatial data can yield more biological insights for complex cellular systems.

## 2 Results

### Overview of VISTA

Our goal is to predict the expression levels of missing genes for all the cells from the spatial data. VISTA jointly models the likelihood of reference scRNA-seq data and target spatial data with geometric machine learning. (Figure 1 (a)). Before fitting our model, we first select reliable genes as anchor genes for these two modalities by thresholding correlation coefficients and significant levels. We assume the gene expression of gene *g* in cell *n* from scRNA-seq data *x*_*ng*_ is sampled from a Zero-inflated Negative Binomial (ZINB) distribution 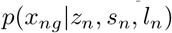 and spatial data expression 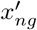 is sampled from a Negative Binomial (NB) distribution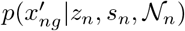. Here *z*_*n*_ represents the joint latent space shared by data from these two protocols, *s*_*n*_ indicates the type of protocol (0 for scRNA-seq and 1 for insitu sequencing). Moreover, for scRNA-seq data, we use *l*_*n*_ to model the sequencing depth, and for spatial data, we use 𝒩_*n*_ to model the spatial information by considering the neighbors of cell *n*. Our target is to approximate the posterior distribution of the latent variables *z*_*n*_, known as *q*(*z*_*n*_, *l*_*n*_ | *x*_*n*_, *s*_*n*_, 𝒩_*n*_), by using a Joint Variational Auto-encoder (JVAE) [34] with Graph Neural Network (GNN). To optimize VISTA, scalable stochastic optimization techniques [35] are employed, ensuring efficiency even with large datasets. Furthermore, the model provides an uncertainty measure for the imputed values based on the distribution of the generative model for each cell and gene (Figure 1 (b)). We sample cells or genes from the distribution and compute the cosine similarity between the sampled data and the estimated mean value for uncertainty estimation. More detailed methodologies of VISTA are described in the Methods section. Our methodology improves both model performances and result interpretation, suggesting the soundness and efficacy of its design.

**Fig. 1.**
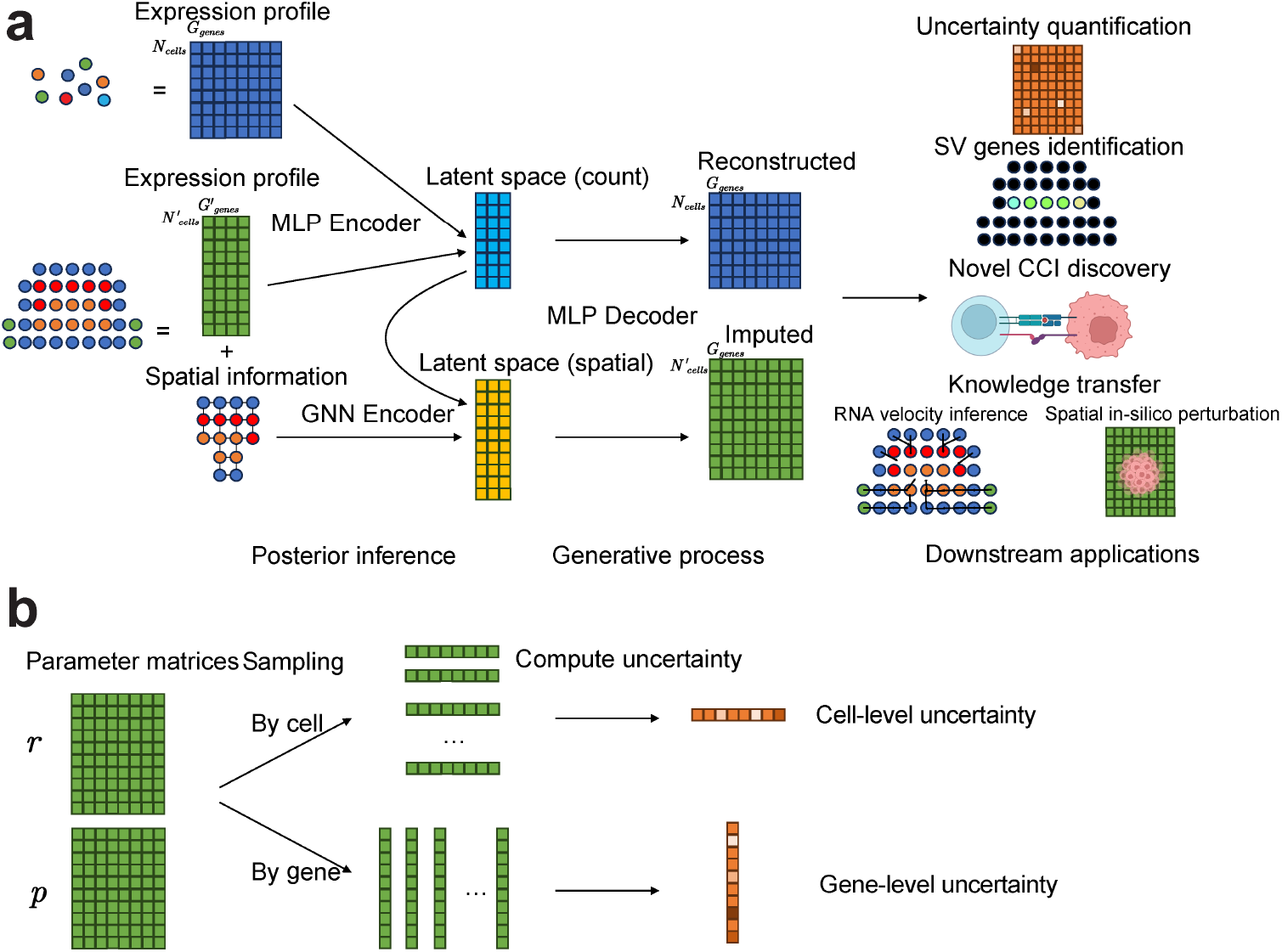
Overview of VISTA. (a): The flowchart of VISTA. VISTA learns a joint distribution from scRNA-seq data and spatial data with graph-aware design. Here we use a Multi-Layer Perceptron (MLP) to encode information from the expression domain and a GNN to encode information from the spatial domain. The results after imputation can be utilized in several downstream analyses. (b): Explanation of uncertainty quantification. The uncertainty quantification of VISTA is based on sampling from the output distribution. For cell uncertainty quantification, we sample from cell-level distribution and compute the median of the cosine similarity between samples and mean gene expression. For gene uncertainty quantification, we sample from gene-level distribution and compute the median of the cosine similarity between samples and mean gene expression.

### VISTA can better impute the expression levels of the unobserved genes with uncertainty quantification than other methods for the osmFISH-brain

#### dataset

Utilizing one dataset known as osmFISH-brain [6, 36], a relatively small-scale dataset commonly used for benchmarking analysis, we demonstrate VISTA’s efficacy. Figure 2 (a) illustrates the distribution of cell types within the spatial data, highlighting specific patterns such as the clustering tendency of Pyramidal cells, hinting at spatial influences on their distribution.

**Fig. 2.**
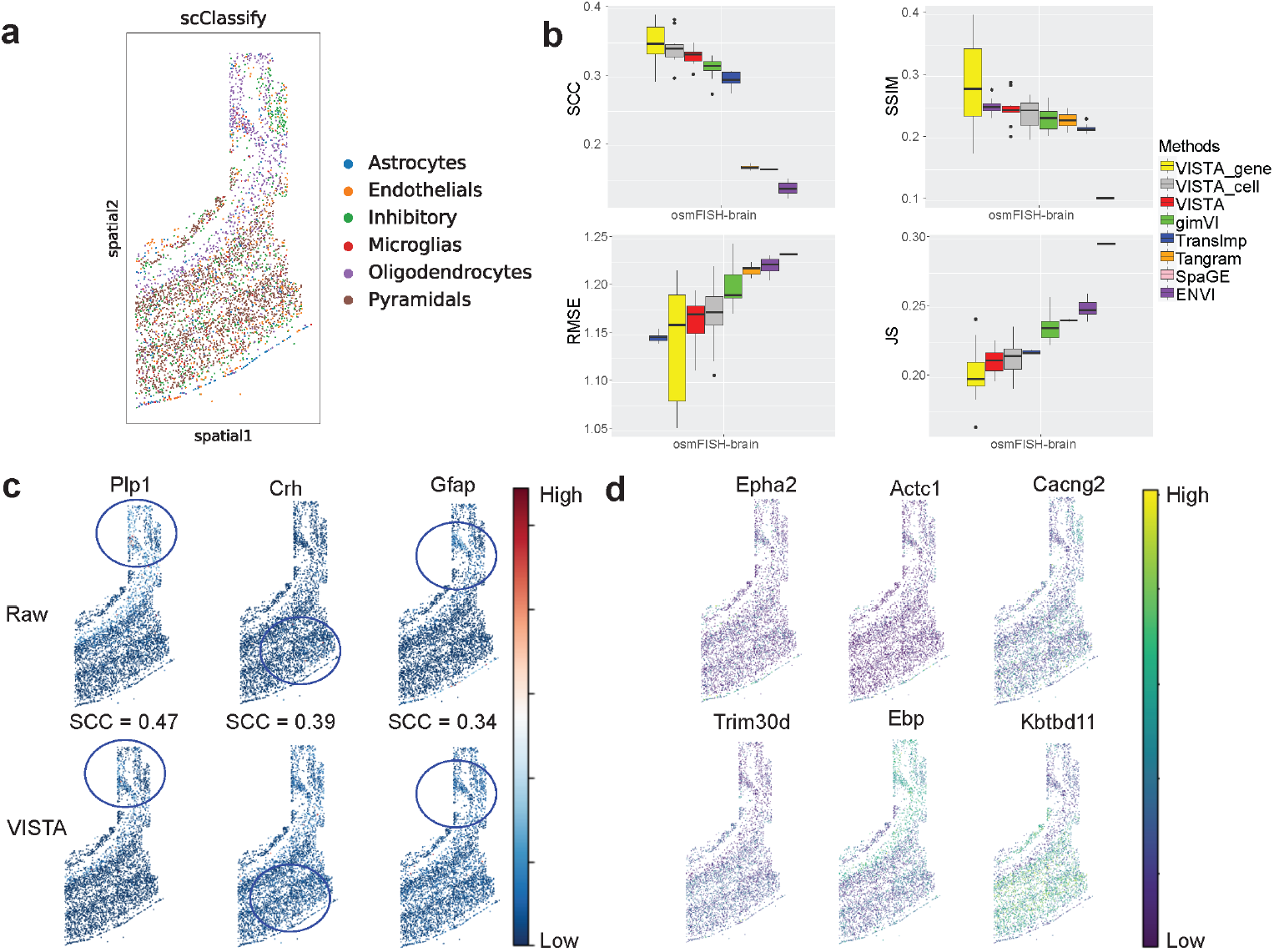
The comparison of imputed results for osmFISH-based data (known as osmFISH-brain). (a) Cell-type distribution across spatial location for osmFISH data. (b) Metrics information of different imputation methods. For SCC and SSIM, higher scores represent better performance, and the methods are sorted in descending. For RMSE and JS, lower scores represent better performance, and the methods are sorted in ascending. (c) Examples of reliably imputed genes by VISTA. (d) Examples of the top marker genes by VISTA for each cell type of osmFISH-brain. The sequence of genes corresponds to the sequence of cells in (a).

The leading three methods for this imputation task are Tangram [24], gimVI [19], and SpaGE [17], as reported by recent benchmarking studies [10, 25]. Moreover, TransImp [28] and ENVI [20] have been introduced as novel tools for imputing spatial data across varying resolutions, and SpatialScope [21] is a method based on diffusion model [37] for spatial solutions in multi-task views including imputation. Consequently, in this study, we consider these six methods for benchmarking analysis, using metrics established in recent benchmarking studies [25]. These metrics were chosen to encompass a broad range of performance dimensions and details can be found in the Methods section. SpatialScope failed in all the datasets due to OOM errors in the diffusion model training step, thus we did not include its results in the following sections. VISTA and other baselines were tuned to their best hyper-parameters, ensuring a fair comparison. As shown in Figure 2 (b), VISTA outperformed other methods in several metrics, such as Spearman correlation coefficient (SCC), structural similarity index measure (SSIM) [38], and Jensen–Shannon divergence (JS), and held a second place in terms of root mean square error (RMSE). These results suggest the better imputation of VISTA.

The comparison and subsequent ablation test further revealed that the integration of spatial information during model training significantly enhances imputation quality. This is evidenced by the superior performance of VISTA over gimVI, which is detailed in Appendix A.

VISTA quantifies uncertainty in two approaches, denoted as VISTA gene and VISTA cell, corresponding to the evaluation of genes and cells with certainty levels above the median, respectively. This quantification process is validated in Figure 2 (b), which shows an increase in SCC when genes or cells with uncertainty rankings in the top 50% are removed. The filtering process, while potentially increasing variance due to a small testing size, still illustrates the reliability of produced results.

The expression patterns of reliable genes, ascertained by VISTA, are shown in Figure 2 (c). The top three genes display a strong alignment between the predicted and observed values, both in terms of distribution in the spatial space and magnitude. In Figure 2 (d), we display the contribution of imputation for discovering novel marker genes of different cell types. The raw marker genes are shown in Extended Data Figure 1. After imputation, we identified more marker genes and linked these genes to the analysis of spatial transcriptomic data. For example, **Kbtbd11** showed strong spatial expression patterns, and cells from Pyramidals also have such patterns. This consistency assures the validity of cell type-related analyses following imputation, emphasizing the role of VISTA in enhancing biological discovery through spatial transcriptomic data refinement.

Finally, we include quantitative comparisons for uncertainty estimation in Appendix A. Again, VISTA performed better than TISSUE in this case.

### VISTA can better impute the Xenium-based and seqFISH-based datasets than other methods

In the context of large-scale spatial transcriptomics, the challenge of imputing missing genes is magnified by the substantial increase in dataset size. The Xenium technique is capable of producing datasets with over ∼ 160,000 cells, whereas the seqFISH technique which can produce ~ 60,000 spots. These techniques offer a stark contrast to the smaller-scale datasets like osmFISH and STARmap. This vast increase in scale presents two main challenges: maintaining accurate imputation while ensuring the scalability to handle such large-scale datasets.

To evaluate imputation methods efficiently for these large Xenium-based [10, 39] and seqFISH-based datasets [40], a bootstrap approach was employed to sample 10% of the cells. This sampling strategy allowed for a practical and computationally feasible benchmark without compromising the robustness of the statistical inference. We can also report variance by bootstrapping data. Figures 3 (a) and (b) show the spatial distribution of cell types within the Xenium-breast and seqFISH-embryo datasets, revealing distinct spatial patterns among certain cell types, such as Malignant cells and Allanois cells, respectively. This visualization supports the hypothesis that spatial pattern is a property of certain cell types. Figure 3 (c) showcases the benchmarking results for the Xenium-breast dataset, with better performance of VISTA to impute unobserved gene expressions within Xenium-based datasets. VISTA ranked at the top for SCC and consistently had high performance across other metrics, together with the additional benefit of uncertainty quantification. Figure 3 (d) shows that VISTA surpasses other methods in terms of the SCC, SSIM, and JS. Notably, when uncertainty-filtered genes are considered, VISTA demonstrates superior performance across all metrics. For the Xenium-breast dataset, Figure 3 (e) shows the genes had high certainty and also had very high SCC, by comparing with the ground truth information for the Xenium-breast dataset. **KRT7** achieved a SCC of 0.9 and the visualization also shows the consistency between the imputed result and the ground truth information. For the seqFISH-embryo dataset, Figure 3 (f) shows that VISTA can also impute the spatial enrichment information of reliable genes. For example, for **Sox4** and **Tcf7l1**, part of the spatial enrichment information of these genes was preserved after imputation.

**Fig. 3.**
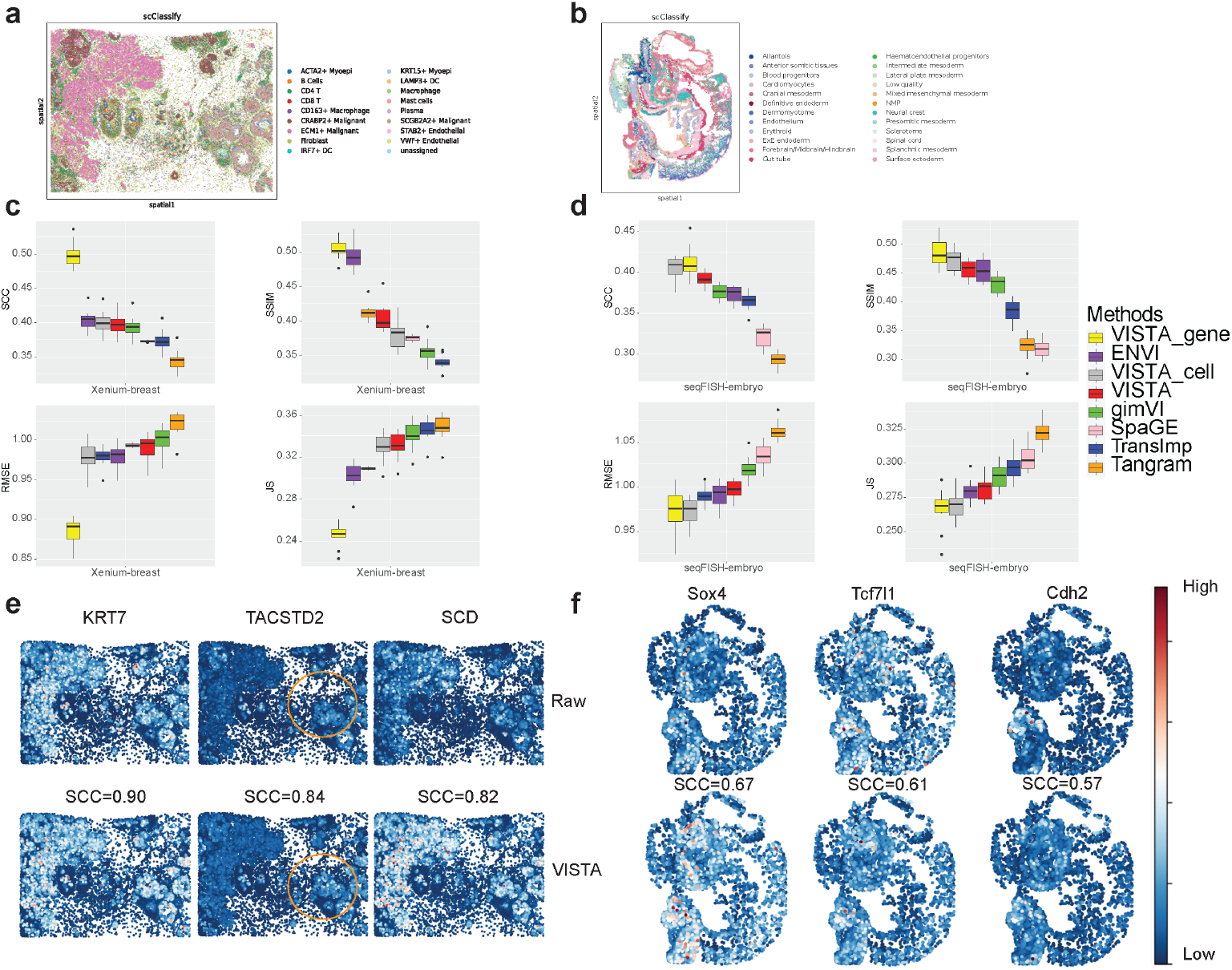
Imputation results comparison for Xenium-based (known as Xenium-breast) and seqFISH-based (known as seqFISH-embryo) datasets. (a) Cell-type distribution by spatial location of Xenium-breast dataset. (b) Cell-type distribution by the spatial location of seqFISH-embryo dataset. (c) Metrics information of different imputation methods for Xenium-breast dataset. (d) Metrics information of different imputation methods for seqFISH dataset. (e) Examples of reliable imputed genes produced by VISTA for Xenium-breast dataset. (f) Examples of reliable imputed genes produced by VISTA for seqFISH-embryo dataset.

### Remarks from evaluation

We discussed the performance of VISTA on imputing the unobserved gene expression levels for the Xenium-brain dataset, which is a Xenium-based dataset, shown in Appendix B.

To compare the performance of different methods across all the datasets (the osmFISH-brain dataset, the Xenium-breast dataset, the Xenium-brain dataset, and the seqFISH-embryo dataset) comprehensively, we averaged the rank of different methods in each dataset and summarized them in Extended Data Figure 2 (a). VISTA outperformed other methods in three out of four datasets by considering the average rank, which implied the strength for imputing missing genes.

### The analysis of uncertainty estimation emphasizes the collection of paired sequencing data

Since the improvement of the uncertainty quantification for the seqFISH-embryo dataset and the Xenium-brain dataset was not clear, we delved into the conditions under which uncertainty quantification can work well as the explainability of VISTA. We hypothesized that the reliability of the imputation results is related to the similarity between the reference scRNA-seq and the target spatial data, with results shown in Extended Data Figures 3 (a)-(c). We used the pre-computed correlations of each shared gene between the reference dataset and the target dataset, plotted the relation between correlations and uncertainty scores, and then computed the SCC based on these two metrics. For the Xenium-breast dataset, we found that the SCC is in negative and significant, which convinced the rationality of our uncertainty quantification method. However, for the results based on the rest two datasets, the correlation was not at a similar significant level (p-value=0.07 for the Xenium-brain dataset and p-value=0.16 for the seqFISH-embryo dataset). Since the reference scRNA-seq dataset and spatial data for the Xenium-breast dataset come from the same tissue, we believed that the imputation of spatial data using a scRNA-seq dataset from the same source yielded more reliable results. Therefore, we advocated the collection of this type of data for analyses.

### The robustness of uncertainty estimation and neighbor searching

Since our uncertainty estimation methods requires the computation of vector similarity as well as the sampling of cells, it is important to demonstrate the robustness and the rationale of current solution with sensitivity analysis. We first considered replacing the cosine similarity (CS) with Euclidean distance (ED) and selected genes with lower ED as genes with higher certainty, and the comparison is shown in Extended Data Figure 4 (a). According to this figure, ED did not perform well in selecting genes with high imputation reliability, which suggests the limitation of using ED to compute distance of high-dimensional data. Furthermore, we also considered alternating the number of sampled cells (*L*) and the result is shown in Extended Data Figure 4 (b). Based on this figure, the results are not sensitive to the choice of L, and thus our uncertainty estimation approach is robust with respect to the tuning.

We also considered adjusting the solutions of neighbor searching based on spatial locations in FAISS [41], including L2 distance (L2), Inner Product (IP), and Locality Sensitive Hashing (LSH), and the results are shown in Extended Data Figure 4 (c). Based on this figure, we also found that using different strategies for neighbor searching did not change our results apparently, and thus our method is also robust to different neighbor searching approaches.

### VISTA facilitates novel discovery of CCIs and SV genes

The process of imputing missing genes in spatial transcriptomics is a critical step for enhancing the resolution of biological functions at the molecular level. One primary goal of this imputation is to enable the identification of novel SV genes and the discovery of new ligand-receptor pairs that are essential for CCI studies, which are central to understanding the complex communications in tissue microenvironments.

Before deploying the imputation models on the entire Xenium-based datasets, a preliminary assessment of the scalability of these models was essential to ensure their feasibility in processing large-scale data without computational constraints. This scalability check is needed to identify any models that may not have been initially designed to handle datasets of such scale, as indicated by the reference to Figure 4 (a). We found that some models including TransImp and gimVI encountered OOM errors when dealing with the extensive Xenium-based datasets (with gimVI failing specifically on the Xenium-brain dataset), VISTA retained its robust performance. Notably, VISTA was not only capable of processing all three datasets but also maintained a relatively stable running time, even as the scale of the datasets increased. This suggests better scalability of VISTA, making it a desirable model for large-scale spatial transcriptomic data. Such efficiency is important, where the volume of data is rapidly growing, and the computational demands are continually escalating.

**Fig. 4.**
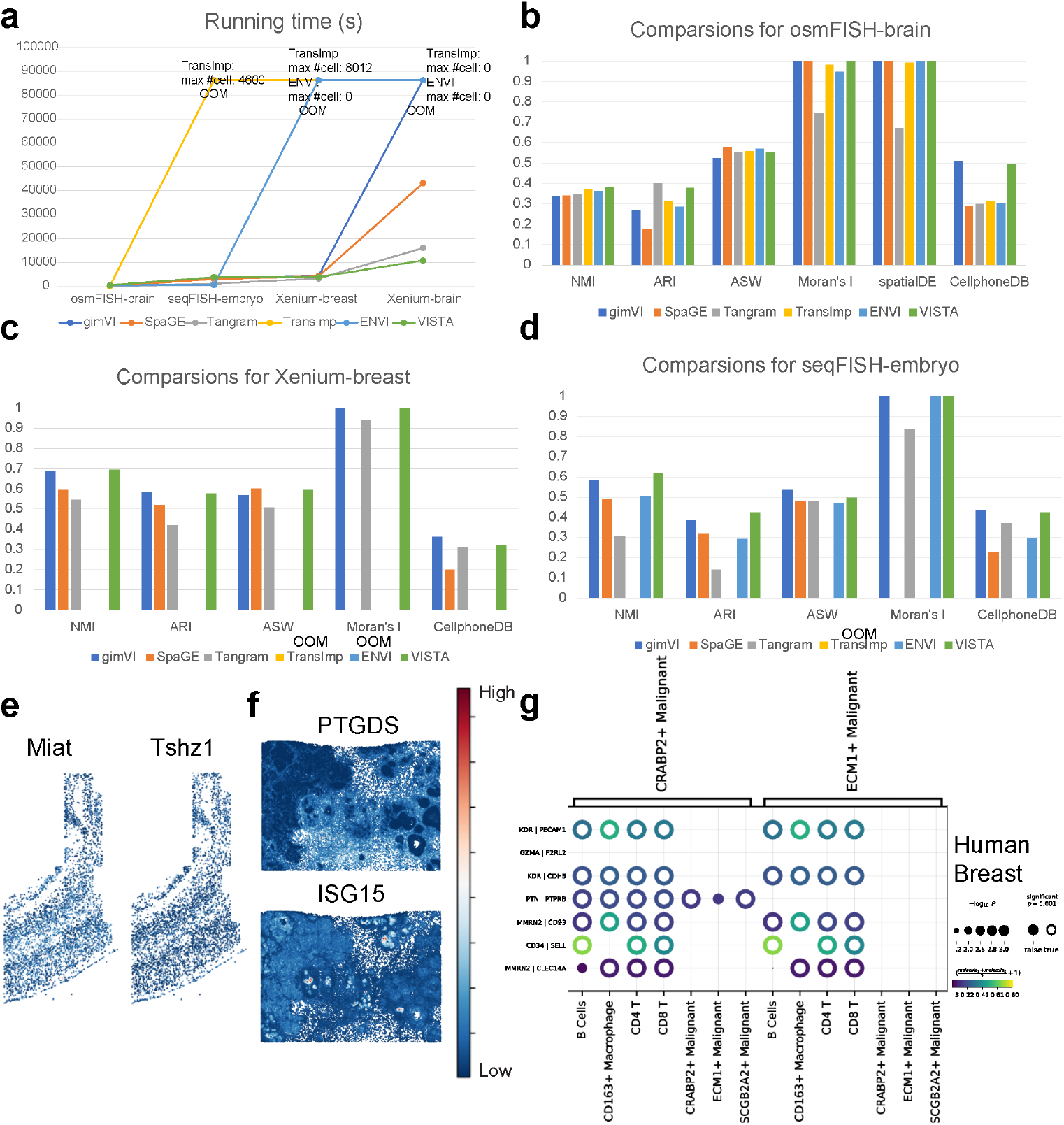
Evaluation of different methods based on scalability and biological function discovery. (a) Running time plot for different methods across the three datasets. We annotate the maximum number of cells for methods that meet OOM errors in this figure. (b) Comparisons of different methods in analyzing the osmFISH-brain dataset. We do not report results from SpatialDE due to out-of-time (OOT) errors and SpatialDE2 due to OOM errors. (c) Comparisons of different methods in analyzing the Xenium-breast dataset. (d) comparisons of different methods in analyzing the seqFISH-embryo dataset. We do not report results from SpatialDE due to OOT errors and SpatialDE2 due to OOM errors. (e) Comparisons of 1st SV gene (left panel) and 1st HV gene (right panel). The plots show the expression distribution of **Miat**. and **Tshz1**. from the imputed osmFISH-brain dataset. (f) Comparisons of 1st SV gene (upper panel) and 1st HV gene (lower panel). The plots show the expression distribution of **PTGDS**. and **ISG15**. (g) Examples of novel ligand-receptor pairs from the results of VISTA, discovered by CellPhoneDB [27] in the imputed Xenium-breast dataset. The color and circle size are explained in the legend part. We take out the parts related to malignant tumor cells and other cells.

Firstly, we considered evaluating the clustering performance of imputation results. We computed the Normalized Mutual Information (NMI), Adjusted Rand Index (ARI) and Average Silhouette Width (ASW) scores [42] based on the clustering after imputation and the original cell-type labels. According to Figure 4 (b) for the osmFISH-brain dataset, VISTA achieved the best NMI score and its ARI and ASW scores were comparable to the best model. Based on Figure 4 (c) for the Xenium-breast dataset, VISTA achieved the best scores for three metrics, and surpassed Tangram and SpaGE. According to Figure 4 (d) for the seqFISH-embryo dataset, VISTA still performed well in the comparison based on the NMI and ARI scores. These results suggest that VISTA is not only robust in imputing missing data but also excels in preserving the biological information of the raw data. The original cell-type-specific information appears to be well-retained after imputation, as reflected in the high scores across different metrics. This fidelity in clustering post-imputation is crucial, as it ensures that subsequent biological interpretations and analyses remain grounded in the cellular heterogeneity before imputation.

Secondly, we considered evaluating the ability of imputation results on SV gene identification. Here we used both Moran’s I score [43] and SpatialDE (as well as SpatialDE2) [26, 44] score to evaluate the correlation between gene expression and spatial location. These two methods have less sensitivity to induced sparsity and SpatialDE has a low false-positive rate [45], and other comparable methods including SPARK-X [46] had OOM errors in analyzing large-scale imputed spatial data. We described these two scores in the Methods section. According to Figures 4 (b)-(d), all methods except Tangram and SpaGE performed well. SpaGE did not find genes whose expression levels were related to spatial location because Moran’s I score was 0. Moreover, VISTA achieved good and stable scores across the three datasets for both Moran’s I score and SpatialDE score. We also compared the imputed gene expression levels of SV genes and Highly Variable (HV) genes. Figure 4 (e) shows that the imputed SV genes discovered by SpatialDE had similar expression patterns with the HV gene identified in the paired scRNA-seq dataset. Due to the running time limit, we recorded the SV gene identified by Moran’s I test rather than SpatialDE and HV gene from paired scRNA-seq dataset for the Xenium-breast dataset, shown in Figure 4 (f), with similar conclusions. Therefore, VISTA’s results incorporate gene variance from paired scRNA-seq data and exhibit similar patterns spatially so we may gain more insights to analyze the similarity among multi-omic data. Overall, these evaluations demonstrate the effectiveness of VISTA in preserving and revealing the spatial heterogeneity of gene expression, which is crucial for understanding the spatial biology of tissues. The stable performance of VISTA across different datasets and metrics demonstrates its utility in the analysis of spatial transcriptomics.

Finally, we evaluated the performance of different methods in discovering novel ligand-receptor pairs. Based on Figure 4 (b), the performance of VISTA was comparable to gimVI and both had promising scores. Based on Figure 4 (c), VISTA ranked 2nd in the comparison. Based on Figure 4 (d), VISTA outperformed Tangram, SpaGE, and TransImp for the seqFISH-embryo dataset. Moreover, Tangram did not perform well across all three datasets, suggesting its limited performance in identifying ligand-receptor pairs. To explore the ligand-receptor pairs discovered in the imputation results by VISTA qualitatively, we display the bubble heatmaps for the Xenium-breast dataset in Figure 4 (g). These identified pairs may piece together complex cellular communication, and lead to a deeper understanding of the functional organization of tissues and potentially highlight targets for therapeutic intervention. For example, we discovered specific highly-expressed pairs KDR-PECAM1 [47, 48], KDR-CDH5 [49], and CD34-SELL [50] between immune cells and malignant tumor cells, thus these genes play an important role for the control of cancers. Overall, the evaluation of VISTA in the context of ligand-receptor pair discovery underscores its potential as a tool for elucidating cell-cell interactions within various biological systems. We also explored the difference between treating the imputed data as scRNA-seq data and treating the imputed data as spatial transcriptomic data, with the results summarized in Appendix C.

Extended Data Figure 2 (b) shows the average rank of metrics we used in this section. VISTA still outperformed the rest of the methods in three out of four datasets. Therefore, the results imputed by VISTA can contribute to various downstream applications.

### VISTA facilitates RNA velocity analysis and signaling direction inference by imputing dynamic properties of genes

Here we explored spatial RNA velocity inference based on spatial transcriptomic data [51, 52]. By replacing the embedding space with spatial location, we can uncover the relation between RNA velocity and spatial distribution by imputing unobserved spliced gene expression and unspliced gene expression. In this section, we analyzed another Xenium-brain dataset sequenced by HybISS [53].

Referencing Figure 5 (a), we conducted a comparative analysis of post-imputation results between TransImp and VISTA, using the RNA data from the scRNA-seq dataset as ground truth. Given that our imputation was derived from the scRNA-seq dataset, we anticipated that the spliced and unspliced RNA profiles in our spatial data would mirror those in the reference dataset. VISTA adeptly conserved the splicing information, whereas TransImp disrupted the expected proportional relationship between spliced and unspliced RNA among different cell types, and inverted the accurate proportion on a global scale. This comparative error analysis is illustrated in Figure 5 (b), suggesting that the imputation of TransImp is unsuitable for inferring RNA velocity. This is further substantiated by the analyses presented on the left of Figure 5 (c), where RNA velocity was computed for both imputation methods using scVelo [54]. The red circle highlights the outcomes from TransImp, where an incongruous convergence of RNA velocity in mature neuronal cells is evident. In contrast, the RNA velocity inferred from VISTA aligns more closely with biological expectations, where mature neural cells are developed from immature ones and there is no clear convergence of RNA velocity in mature cells. Additionally, by employing VeloVI [55], we reassessed RNA velocity to ascertain the level of uncertainty, as depicted in Figure 5 (d). Notably, most cells demonstrated low intrinsic uncertainty versus higher extrinsic uncertainty, indicating that while the magnitude of RNA velocity inferred from imputation is stable, its directional consistency is not.

**Fig. 5.**
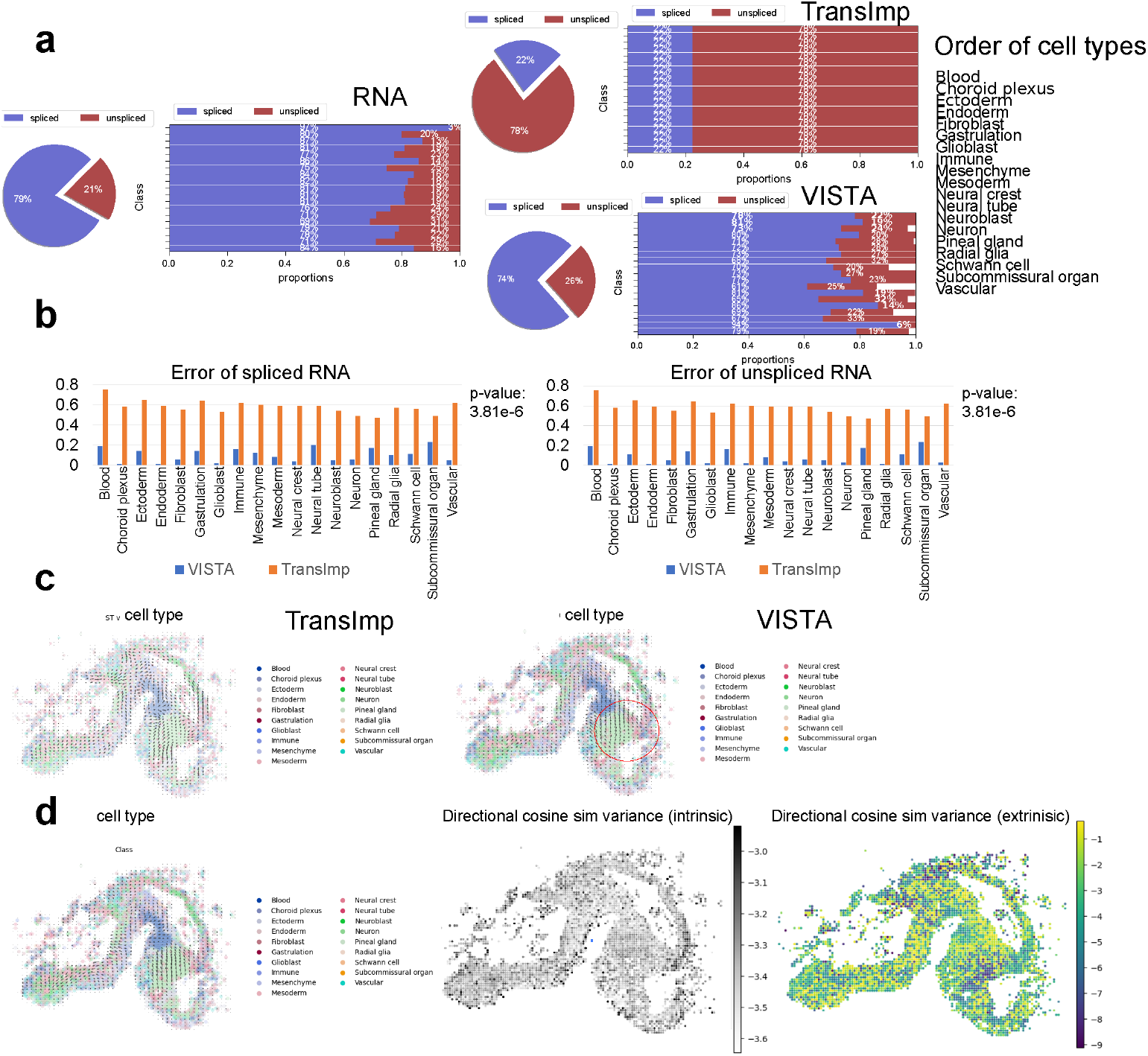
Analysis of spatial RNA velocity inference. (a) Comparisons of spliced/unspliced gene expression proportions. The left panel represents the proportions of the reference scRNA-seq datasets. In the right part, the upper panel represents the proportions of the imputation results based on VISTA, and the bottom panel represents the proportions of the imputation results based on TransImp. (b) Comparisons of the errors between VISTA and TransImp. The errors here are mean absolute errors (MAEs). The left panel represents the errors for spliced RNA and the right panel represents the errors for unspliced RNA. The p-value based on Wilcoxon Rank-sum test of this comparison is also shown in the panel (c) The visualization of RNA velocity based on spatial location using scVelo. The left panel represents the results based on TransImp, and the right panel represents the results based on VISTA. (d) The visualization of RNA velocity based on spatial location using VeloVI. The left panel represents the inference output, the middle panel represents the distribution of intrinsic uncertainty, and the right panel represents the distribution of extrinsic uncertainty.

In addition, we examined the capacity of VISTA to deduce signaling directions within cell-cell communication, a critical aspect of understanding interactions across various cell states. We employed COMMOT [56] to calculate these signaling directions. Figure 6 (a) contrasts the inferred signaling directions from spatial data, both pre- and post-imputation. The left panel of the figure indicates that prior to imputation, the signaling directions computed by COMMOT lacked significance. This was exemplified by the FGF signaling direction, which did not exhibit any discernible pattern in this figure. Post-imputation results, however, were markedly more substantial. The center panel reveals a clear signaling direction for CXCL, as discerned from the imputed data. Similarly, the right panel highlights the ANGPTL signaling direction, bolstering the same inference. Thus, the ability to impute missing gene expressions significantly enhances the detection and analysis of signaling directions, yielding more pronounced and interpretable results.

**Fig. 6.**
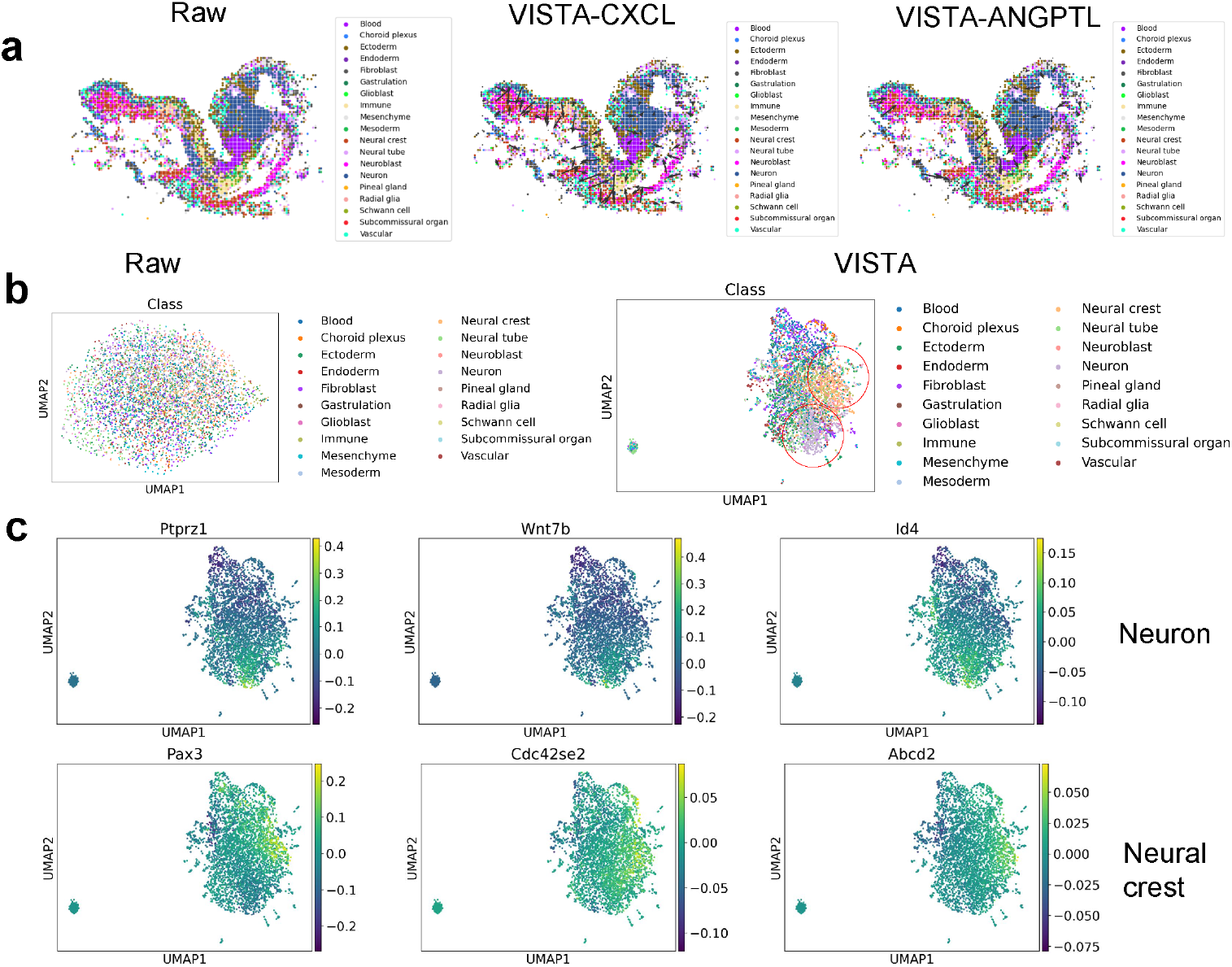
Analysis of signal direction and spatial variation. (a) Results of signal direction based on raw spatial data and imputed spatial data. The left figure represents the signal direction of the FGF pathway based on raw spatial data. The middle figure represents the signal direction of the CXCL pathway based on imputed spatial data. The right figure represents the signal direction of the ANGPTL pathway based on imputed spatial data. (b) Visualization of spatial variation based on UMAP. The left figure represents the spatial variation of raw spatial data and the right figure represents the spatial variation of imputed spatial data. (c) Expression levels of DEGs for cells with spatial-induced cell types. The upper figures represent the DEGs of neuron cells and the bottom figures represent the DEGs of neural crest cells.

### VISTA facilitates the discovery of novel spatial-induced cell types and genes

Decomposing the intrinsic variation (variation brought by cell states) and spatial variation (variation brought by cell-cell interaction at the spatial level) is another pressing open question. Recently, SIMVI [57] was developed to uncover the two different types of variation based on analyzing spatial data. Here we were interested in the difference of these two types of variation by analyzing spatial data before imputation and after imputation. We expected to see that by imputing the expression levels of the unobserved genes, the cells that were affected by spatial factors would be easier to identify. Meanwhile, we could also identify more genes affected by the CCI at the spatial level based on the results after imputation.

To substantiate our proposition, we scrutinized the HybISS and the Xenium-breast datasets to study cell types and genes modulated by spatial factors. We applied SIMVI to both datasets, pre- and post-imputation, and visualized the variance. Constraints in manuscript space led us to present the HybISS dataset analysis within the main text and leave the analysis of the Xenium-breast dataset to Appendix D. From Figure 6 (b), we observed that the spatial data, when not subjected to imputation, failed to produce notable clustering in the UMAPs [58, 59] derived from spatial variance. In contrast, post-imputation spatial data exhibited distinct clustering, particularly for Neuron and Neural crest cells, as indicated by the two red circles. These findings align with the spatial distribution patterns, as shown in Figures 5 (c) and 6 (a).

Additionally, according to Extended Data Figure 5, VISTA demonstrated proficiency in conserving a portion of the intrinsic variance inherent to different cell types. Further analysis revealed the top three differentially expressed genes (DEGs) for these cell types, as determined by the Wilcoxon rank-sum test [60], depicted in Figure 6 (c). These DEGs also displayed prominent signals within the spatial variation-driven clusters. These results support our hypothesis: VISTA can enhance the identification of cell types and genes influenced by spatial factors.

### VISTA facilitates knowledge transfer from single-cell scope to spatial scope

In this section, we extended the capabilities of VISTA to include the transfer of perturbation effects from scRNA-seq to spatial datasets. To simulate the perturbation case, we decoded the gene expression values using latent space from spatial data and decoder part for scRNA-seq data. Firstly, we explored the ability of VISTA to transfer the gene expression levels with diseased information as the special case of perturbation from reference scRNA-seq [61] to target spatial data [62]. We treated the scRNA-seq dataset and the spatial dataset as paired datasets. In total, we have four controlled datasets and four diseased datasets with different ages, thus we separated them into two groups. Here we considered two controlled datasets (one from an 8-month-old mouse, the other from a 13-month-old mouse) and two datasets with Alzheimer’s disease (AD) (one from an 8-month-old mouse, the other from a 13-month-old mouse). We utilized the diseased scRNA-seq dataset as a reference dataset and trained VISTA to transfer the spatial data with the controlled case into the diseased case.

Extended Data Figures 6 and 7 show the results for the 8-month dataset and 13-month dataset, respectively. Extended Data Figures 6 (a) and 7 (a) show the cell-type distribution in the spatial space for AD datasets and control datasets, and we found obvious cell-type overlap between paired datasets. To investigate the correctness of the transferred information based on VISTA, we visualized the overlapping top 10 DEGs from paired datasets with two cases for 8m and 13m in Extended Data Figures 6 (b) and 7 (b). For the overlapped DEGs, the generated AD dataset showed similar patterns compared with the real AD dataset. To perform quantitative comparisons, we computed the differences in gene expression (Δexp) between the spatial dataset with the AD condition and the spatial dataset with the control condition based on Microglia (Micro) cells. Figure 6 (c) shows the Δexp calculated based on the imputed 8m AD dataset and the observed 8m AD dataset. The SCC between these two arrays of Δexp was 0.41 (p-value=1.1e-104). The observation is similar based on Figure 7 (c) for the 13m AD dataset. Therefore, VISTA can successfully simulate the spatial dataset with the AD case. For the other group, we had similar conclusions from Extended Data Figure 8 and Extended Data Figure 9. Moreover, we also considered large-scale spatial data ( ∼ 800,000 cells) [63] sequenced from lung tissue with cancer using MERFISH, as well as large-scale scRNA-seq data ( ∼ 200,000 cells) [64] with diseased information. Extended Data Figure 10 (a) shows the cell-type distribution of the spatial data we used, with clear spatial patterns for Epithelial cells and Fibroblasts cells. After the imputation by VISTA, the specific clustering patterns of these two cells were also preserved, shown in Extended Data Figure 10 (b). Therefore, using the gene expression information after imputation, we could identify more spatial-induced genes for cells from lung cancer samples. The novel DEGs identification after imputation is shown in Extended Data Figure 10 (c). This discovery might contribute to the research of in-silico treatment [65] for lung cancer.

Utilizing a perturbed mouse brain dataset derived from perturb-seq [66] as the scRNA-seq reference, we further assessed the fidelity of transferring perturbation signatures to the previously utilized osmFISH-brain spatial dataset. As depicted in Extended Data Figure 11 (a), the low expression levels of the perturbed gene **Ank2** demonstrate the retention of perturbation effects through our transfer process. Additionally, Extended Data Figure 11 (b) indicates a reduction in the disparity of cell-type distributions post-imputation, suggesting a spatial manifestation of gene perturbation effects. To quantitatively analyze the results we obtained, we selected the genes from the IN1 module, which is affected by the perturbation of gene **Ank2** [66], and computed the correlation between the gene-gene correlation coefficients of the imputed spatial dataset and reference scRNA-seq dataset. In Figure 11 (c), we visualize the correlation with their SCCs. We showed that our imputation process may transfer the perturbed information from reference scRNA-seq data into target spatial data. Further analysis of expression changes in genes common to both datasets, illustrated in Extended Data Figures 11 (c) and (d), confirms the impact of perturbations on gene expression patterns.

Moreover, for scRNA-seq datasets with spliced and unspliced RNA information, we could simulate the spatial transcriptomic data with perturbation by using dynamo [67]. We imputed the corresponding spatial data with both spliced gene expression and unspliced gene expression. We then computed the in-silico perturbation using the RNA velocity from spatial data. We illustrate the dynamic relation of different neural cell types in Extended Data Figure 12 (a), and the dynamic relation corresponded to the differential stages of cells [53, 68–70] showed strong signals. Therefore, we had biological support for the in-silico perturbation inference. We show the perturbation results by suppressing **Gata1** in Extended Data Figure 12 (b) and **Spi1** in Extended Data Figure 12 (c). From these figures, we found that the trajectory was changed in this process and detected fewer trajectory patterns. The activation of these two genes plays an important role in the development of the nervous system [71, 72], suggesting the potential utility of our computation results. Since there are no available datasets for spatial transcriptomic sequencing with perturbation [73] as far as we know, our experiments posed two novel approaches to simulate spatial data under different perturbations for possible downstream analysis [74].

### Final remarks

These insights indicate the potential of VISTA to emulate in-silico perturbation experiments as well as in-silico treatment experiments in spatial data.

## 3 Discussion

Spatially resolved transcriptomics at the single-cell level afford the exploration of cell types and gene patterns across various spatial scales. Yet, the breadth of such analysis was previously constrained by the number of analyzable genes and the cell capture capacity. With the advent of Xenium, we can concurrently sequence numerous cells, circumventing the cell limitation issue. To tackle the gene limitation hurdle, we introduce a computational tool, termed VISTA, which leverages the synergistic modeling of scRNA-seq and spatial data through VAE and GNN. This framework is adept at reconstructing unobserved gene expression levels within spatial datasets. Additionally, the outputs of VISTA can be harnessed for uncertainty quantification and further downstream analyses. Our extensive experimental validations affirm the robustness of VISTA as an imputation method for in-situ sequencing data.

The existing models discussed here serve as valuable benchmarks for the development of our framework, yet they exhibit limitations when applied to large-scale spatial datasets. SpatialScope cannot handle all of the datasets due to memory issues. gimVI fails to integrate spatial context into the modeling process. Moreover, we encountered OOM issues when processing datasets produced via the Xenium platform using gimVI, because its architecture does not efficiently handle large-scale data [75]. Similarly, the initial design of Tangram for cell-spot matching operates on a full-batch basis, leading to OOM during data imputation. Tangram with mini-batch design focuses on local rather than global optimization, thus not reaching its full potential [76]. Additionally, its batch processing necessitates interim file storage, which could be problematic in environments with restricted storage capabilities. SpaGE, employing a regression model within the principal components space, may not adequately discern complex gene-related spatial patterns due to its reliance on linear transformations. Its performance is further hampered by its preference for CPU computation over GPU, affecting computational efficiency. TransImp demonstrates a comparatively improved ability in imputing gene expression levels. However, it fails in processing large-scale spatial data and lacks the capability for uncertainty quantification. Therefore, in the era of Xenium, two pivotal challenges emerge for the imputation task: achieving accurate imputation and maintaining high computational efficiency.

VISTA is a robust solution capable of overcoming the challenges that its predecessors face. Its unique design allows us to handle both small-scale and large-scale spatial data, offering superior performance in various scenarios, including those that require uncertainty quantification. The scalability of VISTA is enhanced by its compatibility with GPU acceleration and its efficient mini-batch training approach, making it a viable option for a wide range of academic research settings due to its reasonable run times on contemporary hardware. The generative model at the core of VISTA supports uncertainty quantification through sampling techniques. We demonstrated the advantages of our uncertainty quantification method by showing the improvement of imputation metrics after removing genes with higher uncertainty as well as comparing it with other baselines. By leveraging this approach, VISTA not only predicts unobserved gene expression levels but also assesses the confidence of these predictions, providing a measure of reliability that is crucial for downstream analysis. The ability to filter and select genes based on their imputation reliability further underscores the utility of VISTA. Consequently, VISTA represents a significant advancement in the field of in-situ sequencing data analysis. Its dual capability to accurately impute unobserved data and quantify uncertainty makes it an invaluable tool for researchers dealing with diverse datasets, whether they originate from established techniques or are the product of cutting-edge technologies like Xenium. This versatility and performance indicate a substantial step forward in computational frameworks for spatial transcriptomics.

In this manuscript, we have explored potential factors influencing model performance, drawing conclusions on how to optimize its use across various datasets. Our findings indicate that incorporating scRNA-seq data along with spatial data from identical tissue and sample sources enhances performance and explainability, as evidenced by analyses using the Xenium-breast dataset. Additionally, there is a need for dataset-specific tuning of VISTA’s hyper-parameters, necessitating user adjustments to attain optimal results. Furthermore, we observed that genes are not imputed equal: Precluding genes with greater uncertainty may enhance result reliability. Lastly, incorporating cell-type information within the training model yielded no substantial improvements. These insights should provide valuable guidance users of VISTA.

Our investigation extended to various downstream applications utilizing imputed results. Firstly, we identified novel marker genes and DEGs after imputation for spatial data with cell-type annotation. Secondly, we discovered genes with spatially expressed patterns by finding SV genes. We also discovered novel ligand-receptor pairs after imputation. Thirdly, we demonstrated the power of VISTA for imputing spliced genes and unspliced genes for spatial data and used the imputation results to infer spatial RNA velocity. Moreover, we also analyzed the signal direction of some pathways to figure out significant patterns related to cell-cell communication. We also discussed the influence of imputation on variation decomposition for spatial data and discovered genes with more significant associations with spatial variation. Finally, we explored possible knowledge transfer from the scRNA-seq domain to the spatial domain. Therefore, the imputation through VISTA offers novel biological discoveries in different dimensions.

Our flexible input feature design enables the imputation of additional modalities, such as Antibody-derived Tags (ADTs) [77] and chromatin accessibility data from single-cell ATAC sequencing (scATAC-seq) [78]. The versatility of VISTA is further illustrated by its applicability to different datasets of similar tissues, a process facilitated by transfer learning [79]. Additionally, the scalability of VISTA positions it as a valuable tool in the construction of spatial cell atlases at single-cell resolution. The potential applications of VISTA are indeed broad and hold great promise for future explorations.

Although we have demonstrated the superiority of VISTA with various experiments, there is room for further improvement. For example, the training of VISTA based on atlas-level data is not stable and requires methods such as gradient clipping to reduce large gradients, and we also did not fully discuss the criteria to select a good reference scRNA-seq dataset for spatial transcriptomics data with different resolutions. In the future, we will consider developing a more unified tool to perform imputation.

## 4 Methods

**Table 1.** STAR Method Table.

| REAGENT or RESOURCE | SOURCE | IDENTIFIER |
| --- | --- | --- |
| **Software and algorithms** |  |  |
| Python 3.8.18 |  | https://www.python.org/downloads/release/python-3818/ |
| PyTorch 1.13.0+cu117 |  | https://pytorch.org/get-started/previous-versions/ |
| scvi-tools 0.20.3 |  | https://scvi-tools.org/ |
| Tangram latest | [24] | https://github.com/broadinstitute/Tangram |
| TransImp latest | [28] | https://transpa.readthedocs.io/en/latest/ |
| ENVI latest | [20] | https://github.com/dpeerlab/ENVI |
| scVelo latest | [54] | https://github.com/theislab/scvelo |
| Dynamo latest | [67] | https://github.com/aristoteleo/dynamo-release |
| **Deposited data** |  |  |
| Codes for reproducing results |  | https://github.com/HelloWorldLTY/VISTA_reproduce/tree/main |

### Problem definition

Considering we have gene expression profiles *X*^*N*,*G*^ and 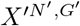 to represent the data from scRNA-seq protocols and the data from spatial transcriptomic protocols, where these two profiles came from the same tissue and *G*^′^ is a subset of *G*. Our target is to learn a model ℳ , which can impute the unobserved genes in spatial transcriptomic data as: 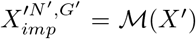. We are also interested in estimating the corresponding uncertainty levels for each cell and each gene. The related downstream applications after imputation are explained later in the Methods section.

### Anchor genes pre-processing

Since we observed that not all the shared genes from scRNA-seq data and SST data are strongly correlated, and some may even have a negative PCC based on Extended Data Figures 13 (a)-(c), a filtering process is needed for both performance and reliability improvement. Here we consider two vectors representing shared cell-type labels as *c*^*M*,1^ from these two datasets. *M* represents the number of shared cell-type labels. We average cells from the same cell types by measurement protocols to generate the pseudo-bulk samples *X*^*M*,*G*^ and 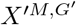. By assuming *G*^′^ ⊂ *G*, we take the common genes and compute the PCC for each gene *g* ∈ *G*^′^, to extract the correlation and significance for each gene:

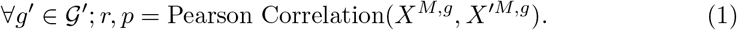

The output variable *r* represents correlation and *p* represents p value. We then filter genes with *r* ≤0.5 or *p* ≥ 0.05 by our default design. The remaining genes are treated as anchor genes and used for constructing the new *G*^′^. We also tested the performances of Spearman correlation but we did not recommend it because the Spearman correlation coefficients of genes are almost all non-significant (p-value*>*0.05), which substantially decreased the size of training dataset and introduced false-negative bias. If users do not intend to include cell types for both reference scRNA-seq data and SST data, they can skip this step.

### Generative model of VISTA

The design of this section is inspired by [19] as a Variational Auto-encoder (VAE) [80]. We assume the following generative process to model the common components of gene expression of scRNA-seq as *x*_*mg*_ and the gene expression of spatial transcriptomic data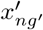:

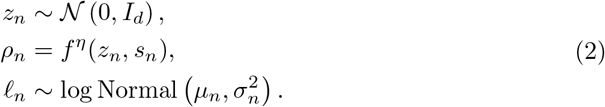

Here we use *z*_*n*_ to represent the shared biology component between the two protocols, and *I*_*d*_ represents an identity matrix with size (*d, d*). For each cell *n*, we have a binary variable as *s*_*n*_ to specify whether the cell is captured by scRNA-seq or spatial experimental protocols. Let *G* donate the set of genes captured by scRNA-seq, and *G*^′^ donate the set of genes captured by spatial transcriptomic data. We assume *G*^′^ ⊂ *G*. We denote a neural network *f* ^*η*^ with parameters *η* to model the probability simplex by generating a variable *ρ*_*n*_. Such variables can be interpreted as the normalized frequencies of each gene *g* of one cell. 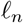 is used to model the log-library size of scRNA-seq data with mean *µ* and variance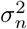 . Correspondingly, we have 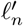 to model the log-library size of spatial transcriptomic data. Since the technical variation of spatial data is not large, 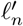 is not a random variable but the number of original gene transcripts in a given cell.

To model the gene expression from scRNA-seq measurement, we treat the observed gene expression for each cell is conditioned on variable set {*l*_*n*_, *z*_*n*_, *s*_*n*_} and the gene expression can be sampled based on overdispersed count data distributions:

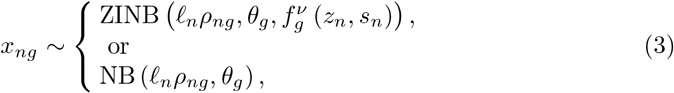

where ZINB represents zero-inflated negative binomial distribution, and NB represents negative binomial distribution. *θ*_*g*_ represents a vector of gene-specific inverse dispersion parameters. Such options are provided by gimVI and we model the gene expression data by ZINB in this manuscript.

To model the gene expression from spatial experimental protocols, we first renormalize the expressed gene frequencies based on:

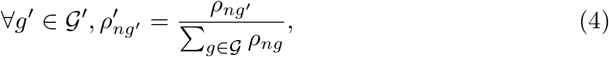

because we subset the genes from *G*^′^. Therefore, we can model the gene expression 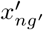_*′*_ also based on 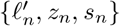 as:

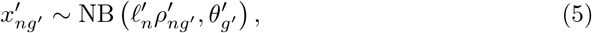

where *θ*_*g*_ represents a vector of gene-specific inverse dispersion parameters. Such design follows the modeling process of gimVI.

### Posterior inference and graph neural network construction of VISTA

The design of this section is inspired by [19], and we utilize a Bayesian inference approach conditioned on spot neighbors to optimize the parameters in the given model. During the training process, to handle the large-scale datasets, we implement an approach based on node sampling to generate neighborhood graphs for each cell in the given batch. For a batch with batch size *n*, we sample *n* cells from the total spatial data and compute the neighbor graph based on these cells. That is:

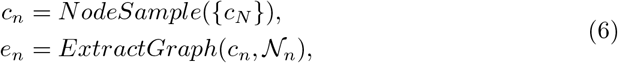

where 𝒩_*n*_ represents the neighbor sets of all the cells in *c*_*n*_. We implement the edge generation process based on FAISS [41] for acceleration.

For the data from two different protocols, we can decompose the objective function as:

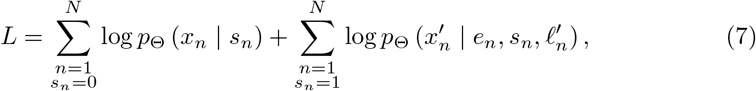

where *N* represents the total number of samples, and *p*_Θ_ represents the distribution under different conditions. The parameters we intend to optimize in the generative model are represented as Θ = {*η, ν, θ, θ*^′^ }. Since it is hard to perform exact posterior inference here, we utilize variational inference to access the posterior distribution.

Considering we have two distributions 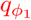 based on scRNA-seq data and 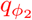 based on spatial data. We can first encode the cell-level embeddings *z* based on a non-linear MLP 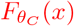 (*x*), and thus we have:

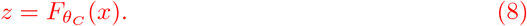

Furthermore, we intend to learn the joint latent space for these two modalities and for data from spatial transcriptomics, so we also connect a Graph Attention Network (GAT) [81] to capture the neighbor information for spatial transcriptomic data. Considering *i* and *j* are two indices of neighbor spots in the training batch, the computation can be represented as:

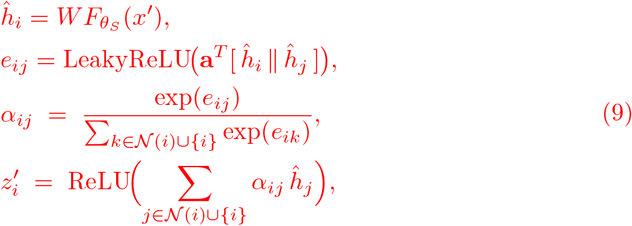

where 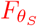 is a non-linear MLP to learn the initial embeddings, *W* and **a** are learnable parameters, LeakyReLU, ReLU are the default activation functions for GAT, *exp* is the function of exponential computing, and 𝒩 (*i*) represents the set of *i*_*th*_ spot’s neighbors. The final updated *z*^′^ represents the spot-level embeddings with spatial context.

Therefore, to formalize the joint distribution, we have:

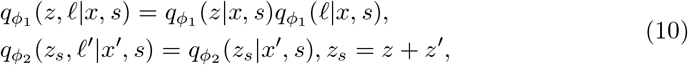

where these distributions are Gaussian with diagonal covariance matrices, and we can utilize a neural network to generate the parameters of these two distributions, which has been discussed above. Here *e* represents the edges of the neighbor graphs of samples in *z*. This setting can also be treated as a version of the mixture-of-expert (MoE) design with one expert focusing on scRNA-seq and the other expert focusing on spatial transcriptomic data [82]. The evidence lowerbound (ELBO) of these two modalities are:

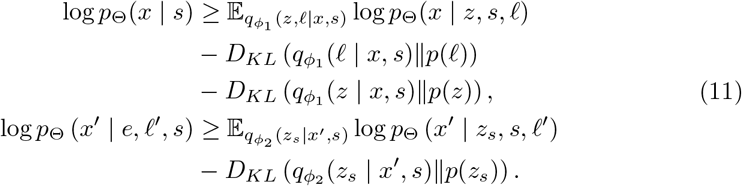

We then optimize the neural networks to obtain the best model, and the aggregation between the latent space as input and the output of GAT is inspired by residual learning [83]. From the domain adaption theory given by [84], we also refer to the implementation of gimVI and add an adversarial classifier for reducing the distribution shift between spatial data and scRNA-seq data. Reliably imputed genes should be either well fitted by VISTA or well fused in the joint latent space.

For the missing gene imputation step, we consider a gene *g*_*i*_ ∈ *G − G*^′^, which is missing in the spatial measurement. We sample the variable *z* based on the posterior distribution *p*_Θ_(*z* | *x*^′^, *s* = 1) and we use our neural networks to compute the missing gene expression 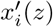 for all the cells in the spatial domain. The mixture model of VISTA is shown in Extended Data Figure 14.

### Uncertainty estimation

We offer the uncertainty estimation at both the cell level and the gene level. Both estimation processes are based on the distribution of the generative model: 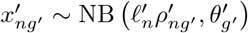 _*′*_ and we denote 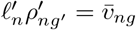, where 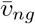 represents the mean of our NB distribution. The estimation process is inspired by VeloVI [55].

For the gene-level uncertainty estimation, we sample *L* examples for all the cells and gene *g*. Therefore, the uncertainty for this gene, known as *δ*_*g*_, can be defined as follows:

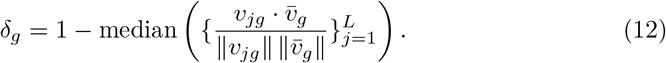

Here *v*_*jg*_ represents the *j*th sample we generate from our output distribution, and we compute the median value of the cosine similarity between the sampled value and mean value. We set 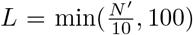 if we have *N* ^′^ cells from spatial data. We use gene-level uncertainty to determine which genes we can trust after imputation, and we select genes whose uncertainty is lower than the median value for comparison in our experiments.

Similarly, for the cell-level uncertainty estimation, we sample *L* cells for cell *c* and all genes. Therefore, the uncertainty of this cell, known as *δ*_*c*_, can be defined as follows:

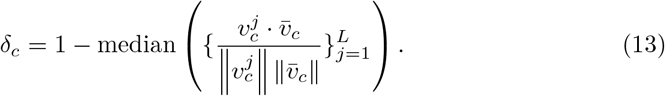

We also set 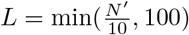 if we have *N* ^′^ cells in spatial data.

### Experiment design

In the benchmarking analysis of imputation accuracy for the four datasets, we split the overlap genes between scRNA-seq datasets and spatial datasets into training datasets (approximately 70% of the genes) and testing datasets (approximately 30% of the genes) [85]. Different methods have their specific design for splitting training datasets into training datasets and validation datasets. To study the stability, we changed the random seeds from 0 to 9 for every method during the experiments. Then we adjust the hyper-parameters of different methods to achieve their best performance in each dataset. The hyper-parameter adjustment information can be found in Appendix A. In the benchmarking analysis of biological functions, we impute all the missing genes of spatial data based on different methods and perform evaluation.

### Baseline models explanation

In our benchmarking process, we consider six baseline methods in total for comparison, including gimVI, Tangram, SpaGE, TransImp, ENVI, and SpatialScope. These methods are randomly ordered. We also included TISSUE for benchmarking the performance of uncertainty estimation.

gimVI [19] also models the gene expression from these two modalities into a joint latent space and uses variational inference to generate the output distribution and impute the expression levels of missing genes. However, gimVI does not consider the neighborhood relation in the spatial data and its implementation is not efficient, as shown in our results. Moreover, gimVI meets OOM errors in our benchmarking process.

Tangram [24] is a model based on optimal matching. Tangram learns the best match relation between scRNA-seq data and spatial data by learning a mapping function and then performs the imputation based on minimizing the loss of such mapping function. However, Tangram is not scalable and the batch version of Tangram does not consider the difference between local optimal solutions and global optimal solutions.

SpaGE [17] is a model based on dimension reduction and regression. It firstly reduces the high dimensions of the input data, and in the joint low-dimensional space, it trains a regression model to impute the value of missing genes. However, SpaGE is not efficient for Xenium-based data with moderate performance.

TransImp [28] is a model based on regression and spatial information regularization. It also relies on dimension reduction for the first step, and in the regression step, it considers both minimizing the loss between predicted data and ground truth data and minimizing the difference of spatial information. However, TransImp meets OOM errors in our benchmarking process and its imputation results lead to poor performance for some downstream applications, for example, RNA velocity inference.

ENVI [20] is a model based on conditional auto-encoder. It learns the embeddings of scRNA-seq data and multiplexed spatial transcriptomic data simultaneously and decodes the embeddings to expression space for imputation. However, ENVI meets OOM issues in our benchmarking process.

SpatialScope [21] is a model based on diffusion model for different tasks related to spatial data. The authors pre-train a generative model based on scRNA-seq data to have a checkpoint. Then they load the model from the checkpoint to fine-tune it based on both scRNA-seq data and spatial data to fill the missing gene expression levels of spatial data based on modelling joint distribution. SpatialScope meets OOM issues in all the datasets we used.

TISSUE [29] is a framework for evaluating the reliability of imputation results. TISSUE treats the training dataset as a calibration dataset and computes the uncertainty measurement score based on the spatial neighbors of the calibration dataset. The final lower and upper prediction errors can be estimated based on the distribution of calibration scores. In our experiments, we only considered the default and recommended version of TISSUE based on SpaGE.

### The contribution of imputation for advancing spatial transcriptomics analysis

By imputing the unmeasured genes’ expression levels based on VISTA, we are capable of gaining more information to better analyze a spatial transcriptomic dataset and make biological discoveries, respectively. In this manuscript, we consider several biological applications which can only be performed after imputation. The first application is the discrimination of cell-type-specific information. Since imputation can help us detect more expressible marker genes for a given cell type, we expect that we are able to better distinguish the characteristics of different cell types through clustering. Secondly, we can discover more cell-cell interactions by performing statistical tests based on more expressed genes, and most of the existing CCIs before imputation can be preserved. Thirdly, observing more expressed genes can also help us better characterize variation contributed by biological signals and spatial signals and decompose them. Finally, imputation allows us to model the dynamic change of gene expression across both time and space. If we utilize a scRNA-seq dataset with spliced and unspliced gene expression information, we can impute a spatial transcriptomic dataset with VISTA and perform RNA velocity analysis based on the given spatial dataset. Moreover, if the scRNA-seq data contain perturbation information, we can also perform in-silico perturbation for a spatial transcriptomic dataset with imputation. This application is a demonstration of VISTA’s transfer learning capacity. The details of the applications discussed above are presented in the following sections.

### Metrics explanations

Our evaluation analysis involves both statistical metrics and biological application metrics. The statistical metrics are based on evaluation for correlation or distribution comparison. The biological application metrics are based on evaluation for the discovery of spatially variable genes and cell-cell interactions.

Inspired by [25], we consider four metrics as statistical metrics, including Spearman correlation coefficients (SCC), structural similarity index measure (SSIM), root mean square error (RMSE), and Jensen–Shannon divergence (JS).

1. SCC. The SCC is defined as:

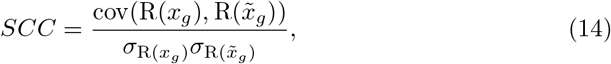

where for gene *g*, we measure the covariance for the rank of ground truth gene expression *x*_*g*_ and predicted gene expression 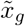 across all the samples. *σ*_∗_ donates the standard deviation for the given ranks. Higher SCC represents better imputation performance, and *SCC* ∈ ( *−* 1, 1).
2. SSIM [38]: Firstly, we scale the gene expression matrix to (0,1) based on:

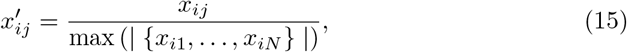

where *x*_*ij*_ donates the gene expression of gene *i* in cell *j*, and *N* represents the total number of cells. SSIM is computed as:

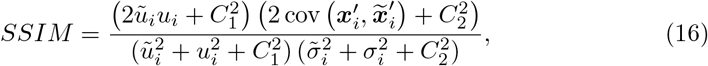

where *u*_*i*_ and 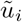 donate the average expression of gene *i* across all the cells. *σ*_*i*_ and 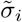 donate the standard deviation of the expression of gene *i* across all the cells. *C*_1_ and *C*_3_ are set a 0.01 and 0.03 based on. Higher SSIM represents better imputation performance and *SSIM* ∈ (0, 1).
3. RMSE. Before computing RMSE, we calculate the normalized score (z-score) for each gene *i* across all the cells, that is:

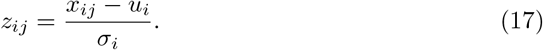 Therefore, for *N* cells, the RMSE is defined as:

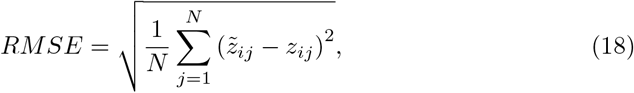

where 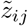 and *z*_*ij*_ denote the predicted z-score and ground truth z-score for gene *i* in cell *j*. Lower RMSE represents better imputation performance, and *RMSE* ∈ (0, ∞).
4. JS. To compute JS, we first define the Kullback–Leibler (KL) divergence for two probability distributions **a**_*i*_ and **b**_*i*_ as:

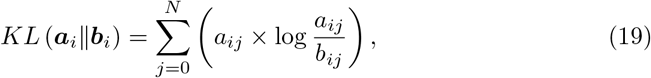

where *a*_*ij*_ and *b*_*ij*_ represent the probabilities of gene *i* in cell *j*. For gene *i*, if we have **P**_*i*_ and 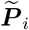 to represent the probability vector of the spatial distribution using ground truth gene expression and predicted gene expression across all the cells, we then can define JS as:

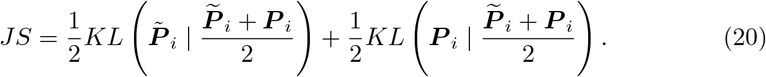

For the evaluation of biological applications, we consider three cases: 1. Evaluating clustering ability; 2. Identifying SV genes; and 3. Identifying cell-cell interaction pairs.

To evaluate the clustering ability, we use three metrics: NMI, ARI, and ASW for evaluation. We choose Louvain [86] as the clustering approach based on the imputation results and treat the original annotation of cell types as ground truth.

1. Normalized Mutual Information (NMI): We calculate NMI score based on computing the mutual information between the optimal Louvain clusters and the known cell-type labels and then take the normalization. Therefore, NMI ∈ (0, 1) and higher NMI means better performance.
2. Adjusted Rand Index (ARI): We calculate ARI score by measuring the agreement between optimal Louvain clusters and cell-type labels. Therefore, ARI ∈ (0, 1) and higher ARI means better performance.
3. Average Silhouette Width (ASW): Here we only consider ASW for cell types. For one cell point, ASW calculates the ratio between the inner cluster distance and the intra cluster distance for this cell. Therefore, higher *ASW*_*cell*_ means better biological information preservation. For *ASW*_*cell*_, we take the normalization, that is:

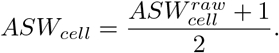

ASW ∈ (0, 1), higher ASW means better performance.

Here we consider two approaches to identify SV genes, known as Moran’s I score [43] and SpatialDE [26] score.

1. Moran’s I score. Moran’s I statistics can test the auto-correlation for spatial data. For one gene, if the corresponding Moran’s I statistics is *I*_*g*_, then we can compute the *z*_*g*_ score by normalization:

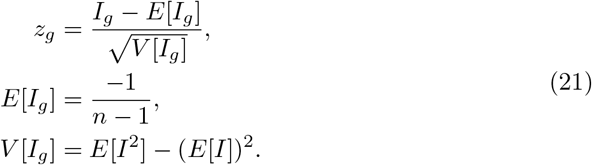 Under the null hypothesis, *z*_*g*_ ∼ *N* (0, 1). We then compute the ratio between the number of genes that are identified as SV genes by setting a threshold based on the p-value and the number of total genes. A higher ratio represents better performance. The scale of the ratio is [0,1].
2. SpatialDE score. For a gene *g* with expression profile *y* = (*y*_1*g*_, …, *y*_*Ng*_) across *N* cells in the spatial domain, we define the given spatial coordinates as *O* = (*o*_1_, …, *o*_*N*_ ). The model of SpatialDE is:

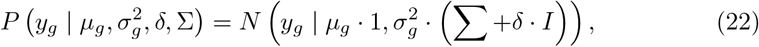

where *µ*_*g*_ and *σ*_*g*_ represent the mean value and standard deviation value for the given gene, and we can determine *δ* based on optimization. The null hypothesis is 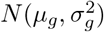, so we can perform the test and get the corresponding ratio after we have all the pvalues. The ratio is defined same as Moran’s I score’s explanation. We also consider SpatialDE2 as a faster version accelerated by GPU cores for its estimation of the Gaussian process. A higher ratio represents better performance. The scale of the ratio is [0,1]. Here we consider three scores to identify cell-cell interaction pairs, which are known as CellPhoneDB [27], CellChat [87] and COMMOT [56].

1. CellPhoneDB score. We use CellPhoneDB to infer the cell-cell interactions for all the cells across cell types, suggested by [88]. CellPhoneDB is based on a permutation test. We first shuffle the cell types for all the cells and take the average of known ligands and receptors in the random cell-type clusters under this case as the null distribution. Correspondingly, we have the observed distribution for the ligand-receptor pairs based on the original cell-type labels. Here also set a p-value threshold for all the discovered ligand-receptor pairs, and compute the ratio between the number of significant pairs and the number of total pairs. A higher ratio represents better performance. The scale of the ratio is [0,1]. The p-value threshold is set as 0.05 for all of the metrics we used in this section.
2. CellChat and COMMOT scores. We also use CellChat and COMMOT to investigate the influence of treating imputed data as different omics to the discovery of CCIs. CellChat and COMMOT both use the CellChat database. CellChat is based on a permutation test for the scRNA-seq dataset, while COMMOT uses the gene expression levels in the whole spots (cells) and distance threshold as criteria for spatial transcriptomic data. Here we use the same ratio we defined in the CellPhoneDB score section. The scale of the ratio is [0,1].

### Application explanation

Besides the biological applications we have mentioned in the metrics explanation section, we also consider three novel applications of spatial data after imputation, including 1. RNA velocity/signaling direction inference, 2. Spatial effect (SE) identification, and 3. In-silico perturbation of spatial data. We do not have ground truth information for these three applications.

For RNA velocity inference, we select scRNA-seq data with spliced and unspliced gene expression as reference data to impute our target spatial data. We impute both spliced gene expression and unspliced gene expression for the spatial data and then run scVelo [54] and VeloVI [55] after imputation. The velocity is computed based on the spatial location.

For the signaling direction inference based on cell-cell communication, we impute the spatial datasets we have and generate the cell-cell signaling direction after imputation based on COMMOT.

For SE identification, we utilize SIMVI [57] to analyze the spatial data before imputation and after imputation. By analyzing the embeddings for spatial effect generated by SIMVI with cell types, we can discover cell types with spatial effect and further genes with spatial effect.

For in-silico perturbation, we consider three conditions. If we intend to simulate spatial data with the diseased condition, we utilize the reference scRNA-seq dataset with the diseased condition as input and generate the corresponding imputed spatial data. If we know the spliced and unspliced information of the reference scRNA-seq data, we use Dynamo to infer the RNA velocity of imputed spatial data and then compute the perturbation under certain genes. If the reference scRNA-seq data has been perturbed without spliced or unspliced information for RNA velocity inference, we treat the raw spatial data as input and utilize the output of the decoder belonging to scRNA-seq data to impute the spatial data with perturbation.

### Data processing

To run VISTA, we keep the raw count data as input since we model the distribution of gene expressions based on the count data matrix.

For the analysis related to biological applications, we have different processing steps. For clustering analysis, we normalized and performed log1p transformation for the raw count data matrix and then selected the top 2000 highly variable genes (HVGs) for the evaluation of the clustering function. For SVG discovery and cell-cell communication discovery, we still use the raw count data. For the analysis of RNA velocity inference, we normalized and performed log1p transformation for the raw count data matrix and then selected the top 2000 HVGs. For the spatial effect identification, we still use the raw count data.

### Datasets information

We did not generate new sequencing datasets in this project. The download links and data statistics are summarized in the supplementary file 1.

### Device and codes information

To run VISTA, we relied on Yale High-performance Computing Center (YCRC) and utilized one NVIDIA A5000 GPU with up to 150 GB RAM. The upper bound of running time for analysis is 24 hours.

The codes of VISTA can be found in https://github.com/HelloWorldLTY/VISTA. The license is MIT license.

## 5 Acknowledgements

We appreciate the comments, feedback, and suggestions from Dr. Jia Zhao for uncertainty explanations, Mingze Dong for the use of SIMVI, and Dr. Romain Lopez for improvement of method description. This project is supported in part by NIH grants R01 GM134005, U24 HG012108.

## 6 Author contributions

Y.L. and T.L. designed this study. T.L. and X.L. designed the model. T.L. ran all the experiments. T.L., Y.L., X.L. and H.Z. wrote the manuscript. Y.S. and H.Z. supervised this work.

## Notes

### Competing Interest Statement

The authors have declared no competing interest.

### Summary of Updates

We have revised the manuscript based on reviewers' comments and provide high-resolution figures for easier reading.

