## Appendix, Supplementary figures, Supplementary file 1 for "VISTA Uncovers Missing Gene Expression and Spatial-induced Information for Spatial Transcriptomic Data Analysis": appendix.pdf

### A Hyper-parameter tuning, ablation tests and sensitivity analysis

**Hyper-parameter tuning.** Here we consider three groups of hyper-parameters for VISTA: epochs, size of latent space, and number of neighbors to construct the graph. The SCC based on different settings of the osmFISH-brain dataset is shown in Extended Data Figures 15 (a)-(d). The optimized hyper-parameters group of VISTA for osmFISH-brain dataset is: epoch=400, n_latent=32, n_neighbors=20, kl_weight = 1.0.

For the seqFISH-embryo dataset, the optimized hyper-parameters group of VISTA is: epoch=200, n_latent=32, n_neighbors=20, kl_weight = 1.0.

For the Xenium-breast dataset and the Xenium-brain dataset, the optimized hyper-parameters group is: epoch=200, n_latent=1024, n_neighbors=20, kl_weight = 1.0.

Note that the choices of the size of latent space and the number of neighbors are also limited by the memory size of our device. When meeting the OOM error, we consider reducing the batch size as a first choice. If it does not work, we consider reducing the size of the latent space followed by reducing the number of neighbors.

For the rest of the methods, the searching space of their hyper-parameters is listed in Table 2.

**Table 2** Hyper-parameter searching scope of different methods. Hyper-parameter settings for SpaGE are from [25].

| Methods | Hyper-parameters |
|---|---|
| VISTA | epochs:[100,500]; n_latent:[10,1024];n_neighbors[5,30] |
| gimVI | epochs:[100,500]; n_latent:[10,1024] |
| Tangram | epochs:[100,500]; |
| SpaGE | adaptive hyper-parameters |
| TransImp | epochs: [1000,3000] |
| ENVI | insensitive to hyper-parameters [20] |

**Ablation tests.** Here we consider six different settings for model design: choices of GNN, choices of graph sampling methods, choices of annotation information, choices of batch-aware design, choices of computing mutual-information, and choices of performing uncertainty estimation. Moreover, we also discuss the importance of filtering low-correlation genes before the imputation process. The SCC based on different settings of the osmFISH-brain dataset is shown in Extended Data Figures 16 (a)-(f). Based on the test results, our final VISTA utilized GAT as a GNN model rather than other candidates [30, 89, 90] to incorporate neighbor information of spatial data because of its high score and efficiency. The final version of VISTA also utilizes node sampling [32] rather than cluster sampling [91] or neighbor sampling [32] for large-scale graph because of performance, and we also avoid including cell-type information [92] in the imputation model design. We do not use the original batch-aware design in gimVI, which is based on modeling the batch information in the dispersion term of the generative distribution, because we found an obvious performance drop if we include scRNA-seq data from multi-sources for imputation. Such drop happens in both the batch-aware case and batch-unaware case. Therefore, incorporating batch labels from different batches with batch effect does not contribute to VISTA. Finally, Extended Data Figure 16 (e) shows the contribution of removing genes with lower correlation and significance levels. Moreover, our filtering process also helps reduce the variance for the imputation results. Finally, we investigated the contribution of latent space disentanglement for imputation. Recent work suggests that minimizing the mutual information between the latent space with graph information and the latent space without graph information can help in fitting VAE and modeling spatial transcriptomic data. Based on Extended Data Figure 16 (f), we found that we could not have better performance by including such design. One reason might be that the shared information between graph-level data and non-graph-level data is both important for spatial transcriptomic data imputation. Based on Extended Data Figure 16 (g), we demonstrated that our uncertainty estimation approach is more reliable than TISSUE, since after filtering the cells and genes with high uncertainty, the SCC from VISTA is always higher than that of TISSUE. We considered genes or cells with uncertainty scores in the bottom 50% to be reliable.

**Sensitivity analysis.** Here we explored the sensitivity of different methods towards the change of the size of training datasets. By using different proportions of cells for both spatial data and scRNA-seq data, we plotted the relation between such proportion and SCC score in Extended Data Figures 17 (a) and (b). From these two figures, we found that all of the current methods are robust to the change of cell proportion. Moreover, VISTA still outperformed Tangram and SpaGE under different proportions, and VISTA performed better than gimVI and TransImp in eight out of nine cases for changing spatial data proportion and seven out of nine cases for changing scRNA-seq data proportion. Therefore, the superiority of VISTA is preserved under the settings of different numbers of cells.

Moreover, by adjusting the size of genes for testing (same as the proportion of genes for testing), we found that increasing the testing size could decrease the performance of the model, shown in Extended Data Figure 17 (c). Such a result implied that in the real application, we should rely on all the overlap genes after filtering as genes for training.

We provided the codes used to reproduce the results of VISTA in https://github.com/HelloWorldLTY/VISTA_reproduce.

### B Extra analysis for the Xenium-brain dataset

In this section, we present an analysis of the imputation performance for the Xenium-brain dataset using various methods, along with the downstream applications enabled by these imputations. The spatial distribution of cell types within the dataset is depicted in Extended Data Figure 18 (a), highlighting discernible spatial patterns, such as the CA1-3 cells. According to the benchmarks in Extended Data Figure 18 (b), the selection of genes with higher certainty allowed the identification of genes with reliable quality, even in cases where VISTA was not the top performer. This underscores the robustness of VISTA in handling the complexities of Xenium-based datasets, both in imputing missing genes and in quantifying uncertainty. Moreover,

Figure 18 (c) shows that VISTA could also impute the spatial enrichment information of reliable genes. For example, for **Neto2** and **Rorb**, part of the spatial enrichment information of these genes was preserved after imputation. As for biological function, as assessed in Extended Data Figure 18 (d), VISTA not only maintains the original cell-type distribution but also identifies additional Spatially Variable (SV) genes and novel ligand-receptor interactions by leveraging a broader gene dataset. Meanwhile, TransImp and gimVI encounter OOM errors when attempting to impute all absent genes. Thus, VISTA proves its efficacy in processing datasets of comparable structure and scale to those based on seqFISH.

### C Extra analysis for discovering CCI

In this section, we discuss whether the results after imputation should be treated in the same way that spatial transcriptomic data is treated for CCI discovery. Extended Data Figure 2 (c) displays the average rank for using CellChat-based method rather than CellPhoneDB to detect CCI across different datasets. From this figure, we found that the performance of VISTA is not outstanding. The results of SPARKX based on the osmFISH-brain dataset is shown in Extended Data Figure 2 (d), and SPARKX is only executable in this dataset. Furthermore, we investigated the detailed scores for these two CCI identification methods, shown in Extended Data Figure 2 (e). The first thing we discovered is the obvious difference in the magnitude of scores based on different methods. CellChat had a similar score compared with the results of CellPhoneDB. Therefore, CellChat tended to select more ligand-receptor pairs, while COMMOT was more reserved. Moreover, for the Xenium-brain dataset, COMMOT did not detect any ligand-receptor pairs based on the results of VISTA, while CellChat only detected ligand-receptor pairs based on the results of VISTA. Such a result implied that VISTA tended to transfer the information from the spatial domain and scRNA-seq domain to a space more similar to the scRNA-seq domain. Such conclusion is supported by the score difference between COMMOT and CellChat and the score similarity between CellChat and CellPhoneDB. Moreover, since there exist both short-range CCI and long-range CCI [88], using distance information or spatial information as a threshold may not be good for long-range CCI detection.

### D Extra analysis for the Xenium-breast dataset

Here we analyze the intrinsic variation and spatial variation based on the Xenium-breast dataset, and the results are shown in Extended Data Figure 19. To run SIMVI based on datasets sequenced by Xenium, we subsampled 10% of the original Xenium-breast dataset for the downstream analysis here. Extended Data Figure 19 (a) shows that the variation caused by cell types is well preserved after imputation. Figure 19 (b) shows that imputation results can uncover the spatial effect of certain cell types, for example, ECM 1+ Malignant and B cells. The spatial variation of these cell types cannot be extracted before imputation, although we can observe the spatial patterns of these cell types. Furthermore, we analyzed the DEGs of these cell types, shown in Figure 19 (d). In this figure, the DEGs of these cell types also showed the specific expression patterns related to spatial variation.

### E Supplementary figures

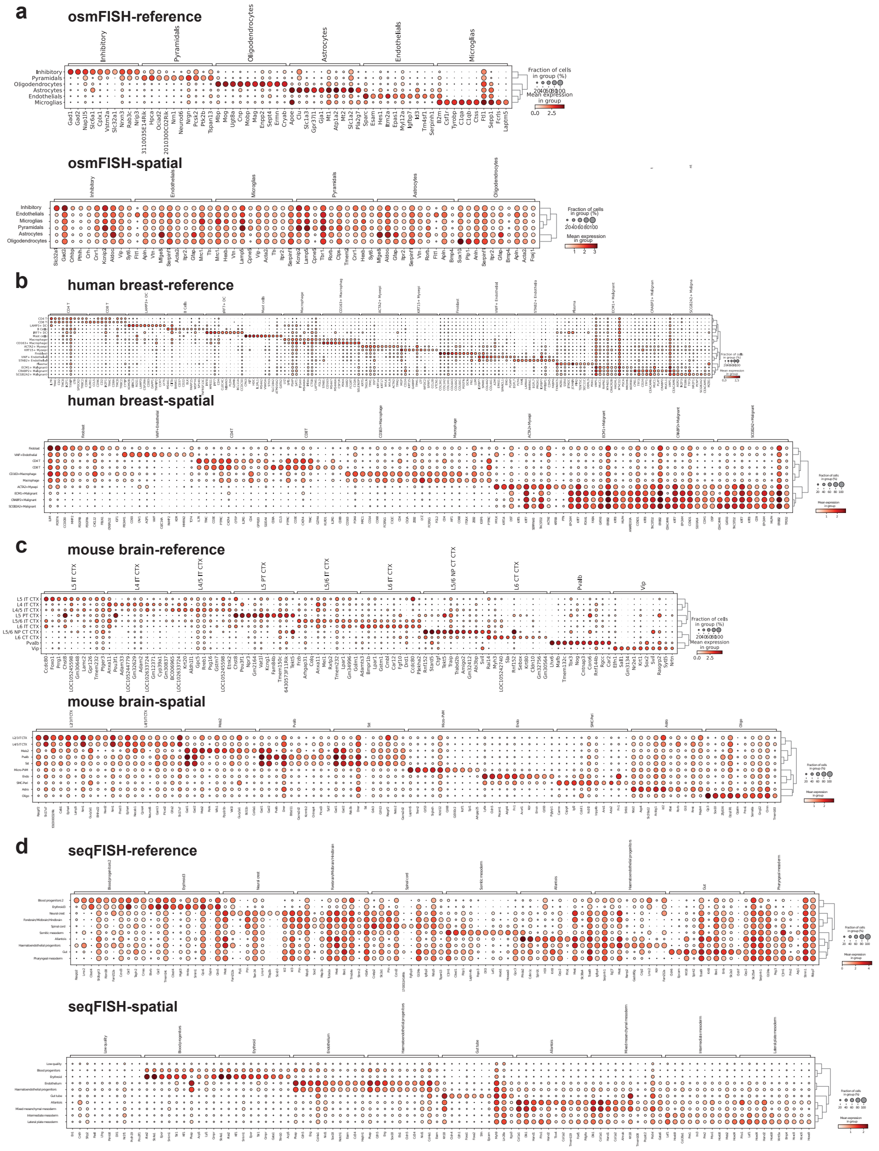


**Extended Data Fig. 1** DEGs of reference scRNA-seq data for different spatial datasets. (a) DEGs from osmFISH-brain dataset of the reference scRNA-seq (upper panel) and the spatial dataset (bottom panel). (b) DEGs from Xenium-breast dataset of the reference scRNA-seq (upper panel) and the Xenium-breast dataset (bottom panel). (c) DEGs from Xenium-brain dataset of the reference scRNA-seq (upper panel) and the Xenium-brain dataset (bottom panel). (d) DEGs from seqFISH-embryo dataset of the reference scRNA-seq (upper panel) and the spatial dataset (bottom panel).

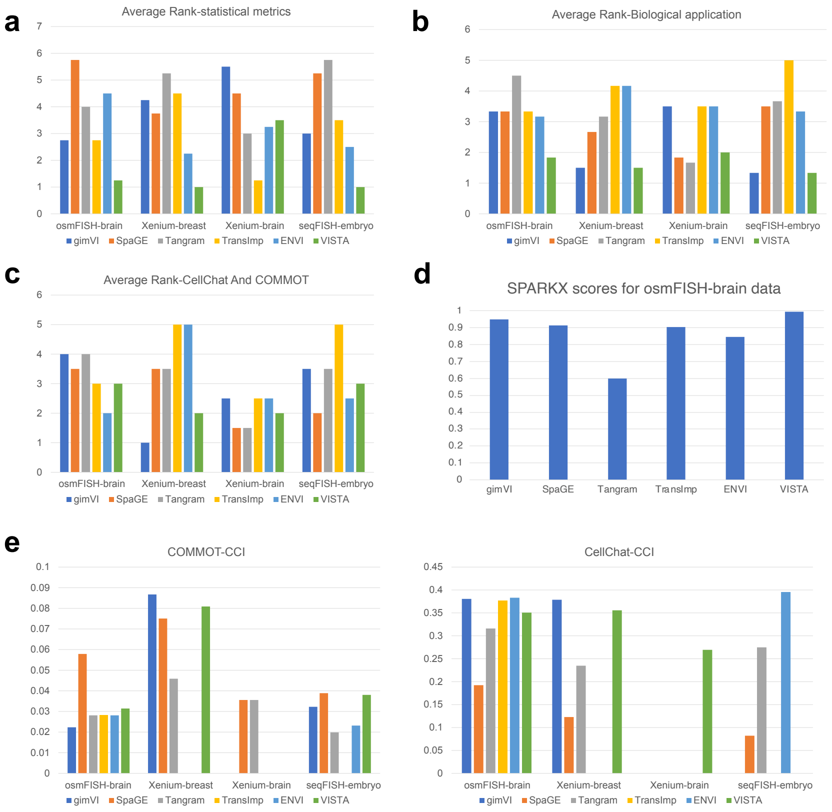


**Extended Data Fig. 2** Average rank and CCI benchmarking score for different datasets. (a) The average rank of statistical metrics across different datasets. (b) The average rank of biological application metrics across different datasets. (c) The average rank of the CCI benchmarking score is based on the CellChat database. CCI score computed based on CellChat and COMMOT. CellChat treats the input data as scRNA-seq data and COMMOT treats the input data as spatial transcriptomic data. These two methods both use the database from CellChat. (d) The significant ratio of different methods for COMMOT (left panel) and CellChat (right panel) across different datasets.

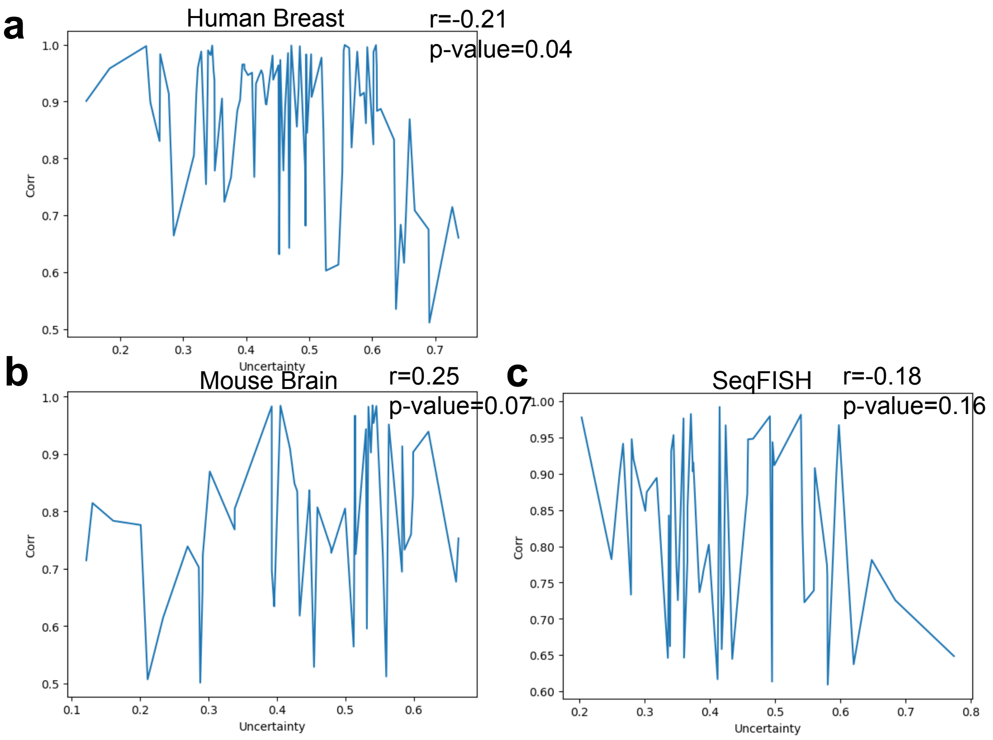


**Extended Data Fig. 3** Visualization for the relation between gene-gene correlation of paired training dataset and uncertainty levels. We used genes in the testing group for computation. (a) Correlation of genes between scRNA-seq and spatial data vs. uncertainty levels based on the Xenium-breast dataset. (b) Correlation of genes between scRNA-seq and spatial data vs. uncertainty levels based on the Xenium-brain dataset. (c) Correlation of genes between scRNA-seq and spatial data vs. uncertainty levels based on the seqFISH-embryo dataset.

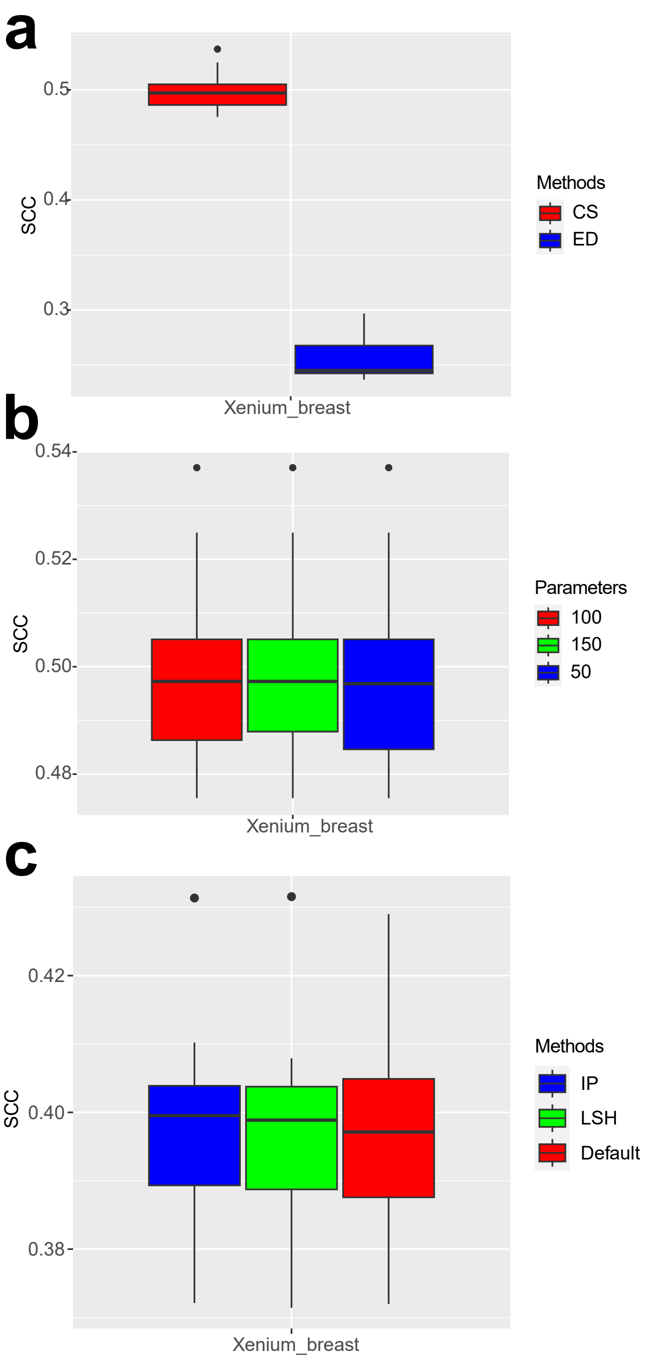


**Extended Data Fig. 4** Sensitivity analysis for uncertainty estimation and neighbor searching. (a) The SCC between imputed and observed gene expression levels based on different approaches for distance computation. (b) The SCC between imputed and observed gene expression levels based on different numbers of sampled cells. (c) The SCC between imputed and observed gene expression levels based on different neighbor searching approaches.

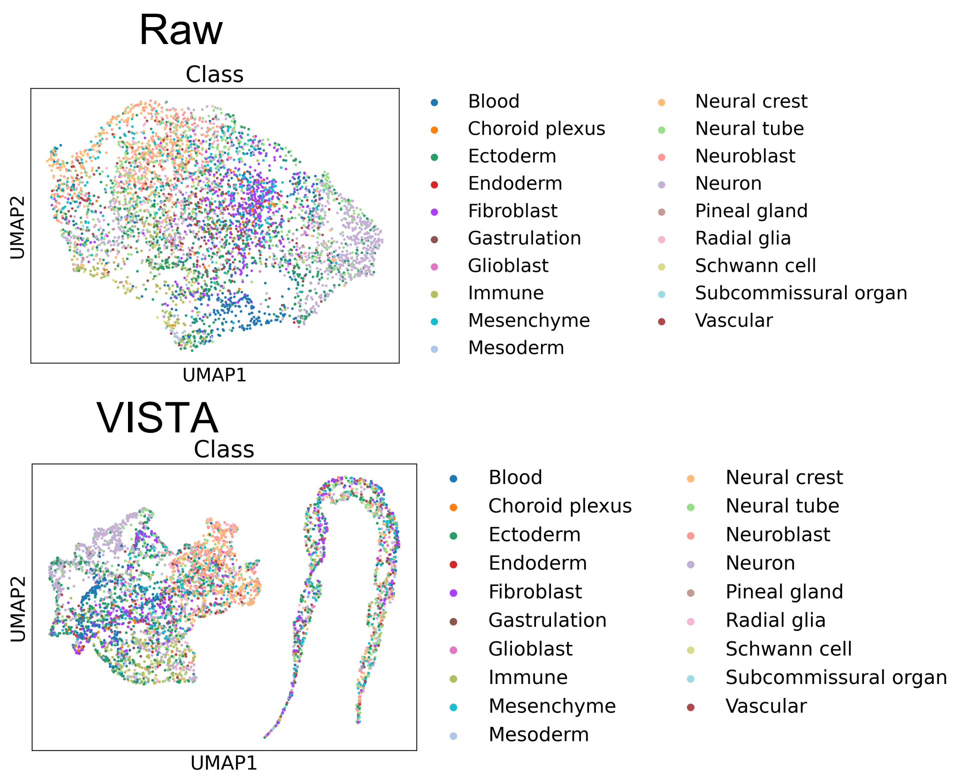


**Extended Data Fig. 5** UMAPs of the intrinsic variation colored by cell types for the raw spatial data and imputed spatial data for the HybISS dataset.

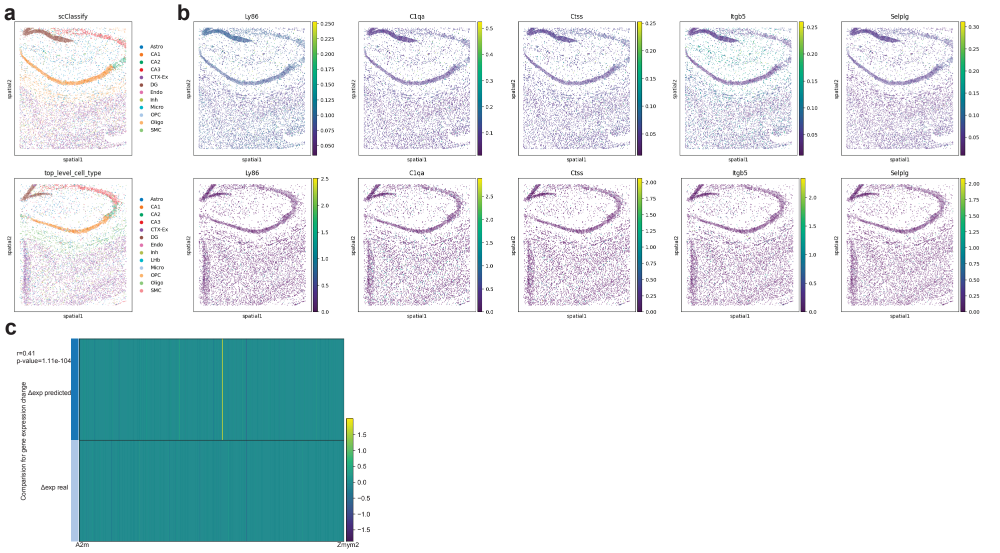


**Extended Data Fig. 6** Results for the in-silico perturbation with 8m dataset (rep1). (a) Cell-type distribution by the spatial location of the 8m mouse brain dataset with control case (upper panel) and dataset with disease case (bottom panel). (b) Gene expression profiles of overlapped HVGs between the transferred dataset (upper panel) and the ground truth 8m mouse dataset with disease case (bottom panel). (c) Visualization of $\Delta exp$ of averaged gene expression levels by cells. $\Delta exp$ *predicted* represents averaged gene expression levels of predicted gene expression minus the averaged gene expression levels of the control dataset. $\Delta exp\ real$ represents the averaged gene expression levels of the AD dataset minus the averaged gene expression levels of the control dataset. We also report the SCC and p-value in this figure.

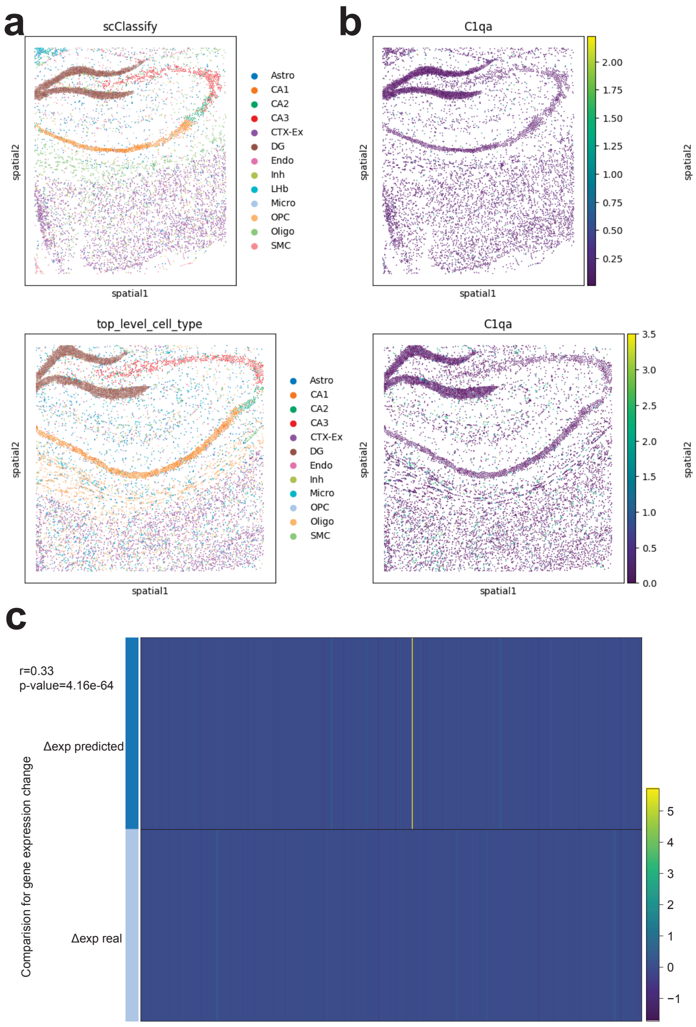


**Extended Data Fig. 7** Results for the in-silico perturbation with 13m dataset (rep1). (a) Cell-type distribution by spatial location of the 13m mouse brain dataset with control case (upper panel) and dataset with disease case (bottom panel). (b) Gene expression profiles of overlapped HVGs between the transferred dataset (upper panel) and the ground truth 8m mouse dataset with disease case (bottom panel). (c) Visualization of $\Delta exp$ of averaged gene expression levels by cells. $\Delta exp$ *predicted* represents averaged gene expression levels of predicted gene expression minus the averaged gene expression levels of the control dataset. $\Delta exp$ *real* represents the averaged gene expression levels of the AD dataset minus the averaged gene expression levels of the control dataset. We also report the SCC and p-value in this figure.

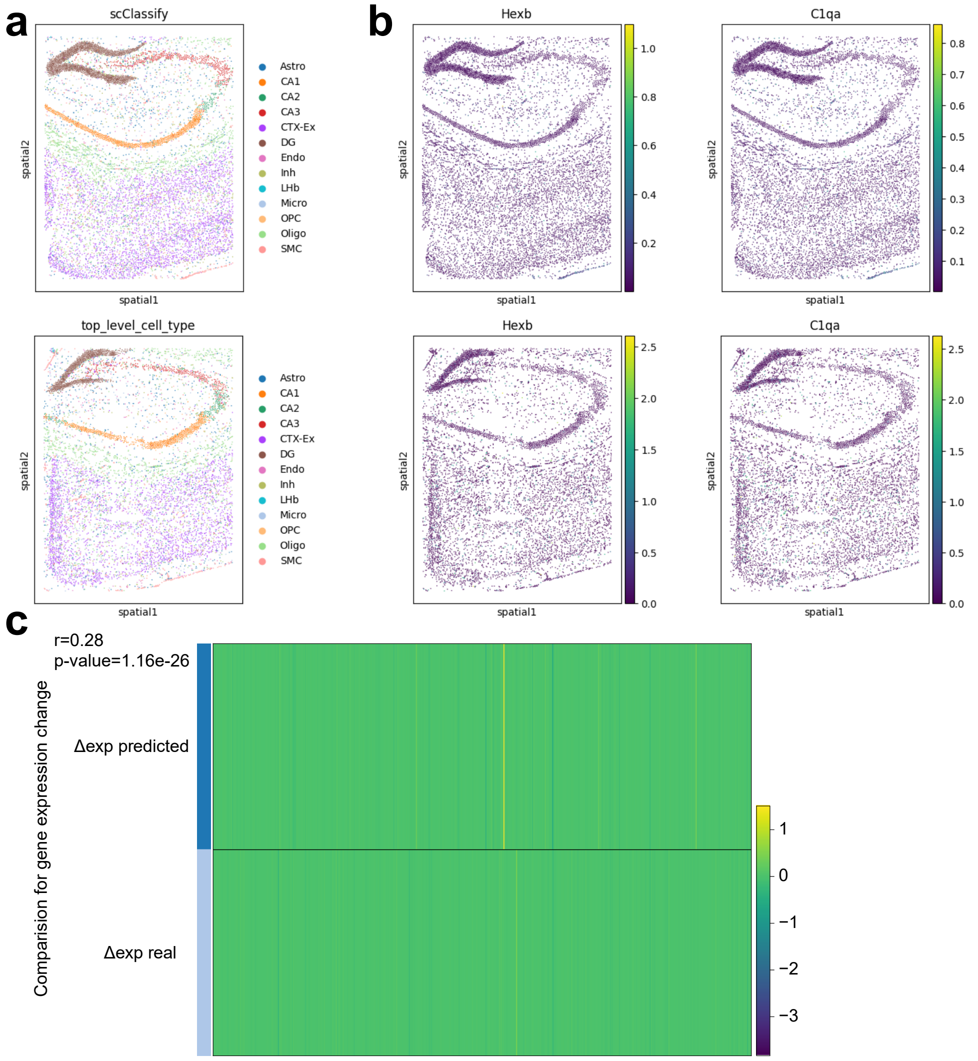


**Extended Data Fig. 8** Results for the in-silico perturbation with 8m dataset (rep2). (a) Cell-type distribution by the spatial location of the 8m mouse brain dataset with control case (upper panel) and dataset with disease case (bottom panel). (b) Gene expression profiles of overlapped HVGs between the transferred dataset (upper panel) and the ground truth 8m mouse dataset with disease case (bottom panel). (c) Visualization of $\Delta exp$ of averaged gene expression levels by cells. $\Delta exp$ *predicted* represents averaged gene expression levels of predicted gene expression minus the averaged gene expression levels of the control dataset. $\Delta exp$ *real* represents the averaged gene expression levels of the AD dataset minus the averaged gene expression levels of the control dataset. We also report the SCC and p-value in this figure.

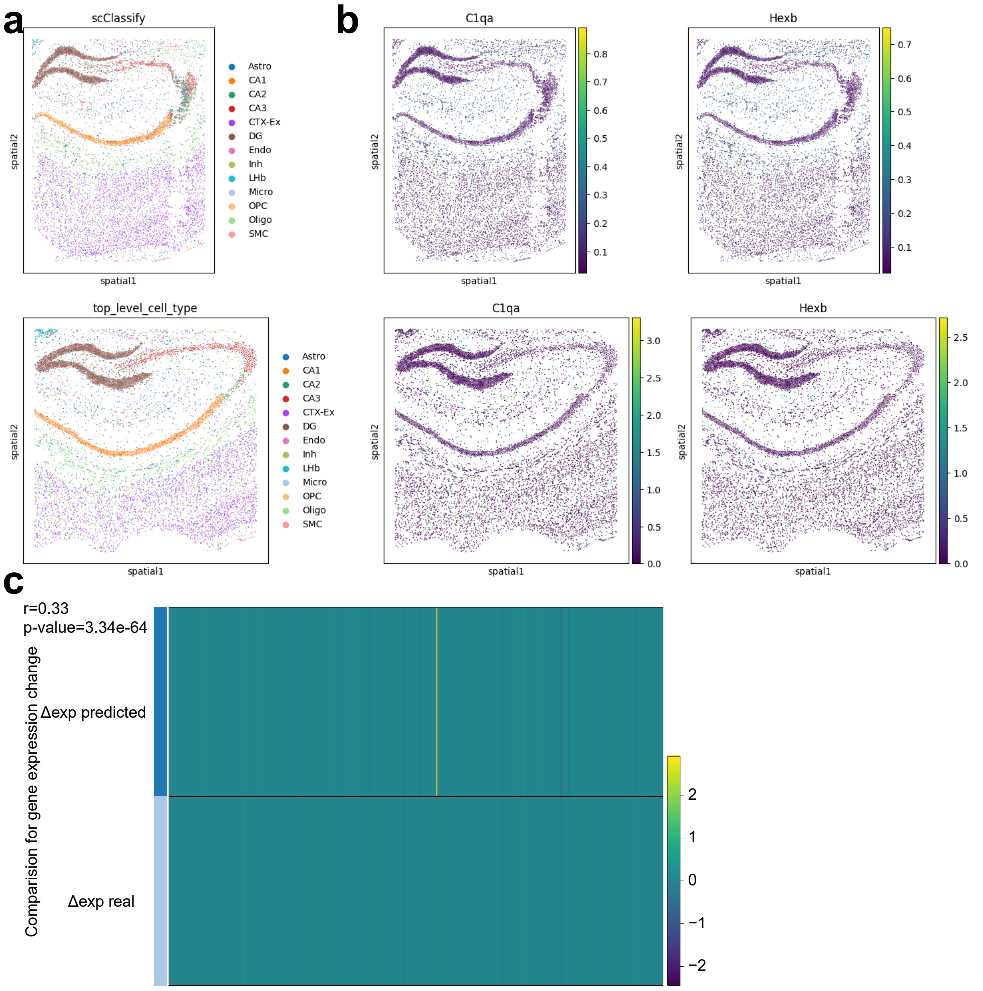


**Extended Data Fig. 9** Results for the in-silico perturbation with 13m dataset (rep2). (a) Cell-type distribution by spatial location of the 13m mouse brain dataset with control case (upper panel) and dataset with disease case (bottom panel). (b) Gene expression profiles of overlapped HVGs between the transferred dataset (upper panel) and the ground truth 8m mouse dataset with disease case (bottom panel). (c) Visualization of $\Delta exp$ of averaged gene expression levels by cells. $\Delta exp\ predicted$ represents averaged gene expression levels of predicted gene expression minus the averaged gene expression levels of the control dataset. $\Delta exp\ real$ represents the averaged gene expression levels of the AD dataset minus the averaged gene expression levels of the control dataset. We also report the SCC and p-value in this figure.

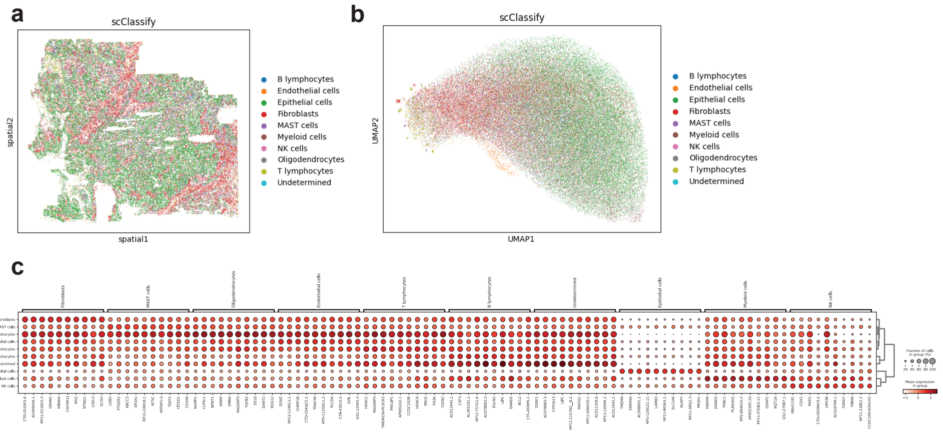


**Extended Data Fig. 10** Analysis of disease information transfer of the human lung cancer dataset. (a) Cell-type distribution by spatial location of the lung cancer dataset. (b) UMAPs of the imputed results by cell types. (c) DEGs of the imputed spatial data by cell types.

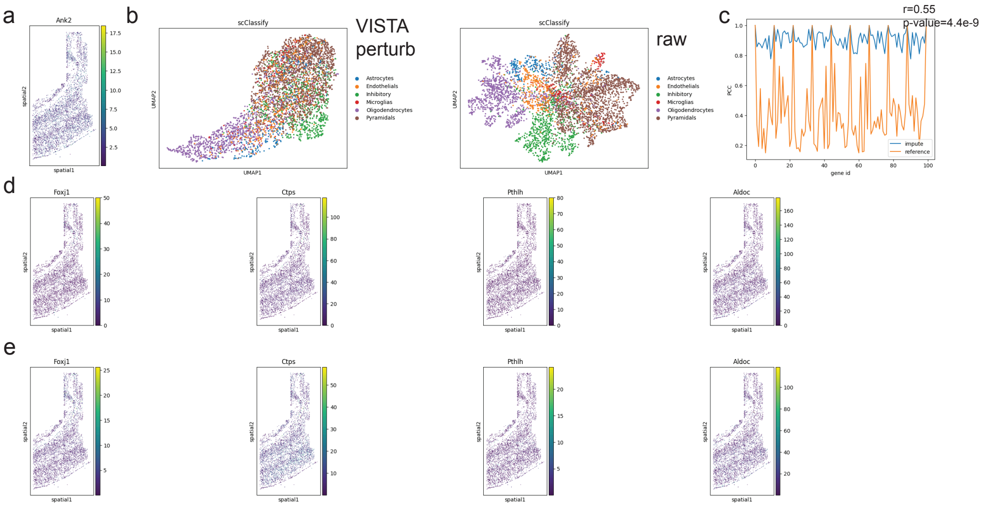


**Extended Data Fig. 11** Analysis of spatial perturbation simulation. (a) Expression levels of the perturbed gene based on spatial dataset after imputation. (b) UMAPs of the cell-type distribution after perturbation (left panel) and before perturbation (right panel). (c) The plot for the relation between gene id and gene-gene correlation coefficient is based on the imputed dataset (impute) and reference dataset (reference). We also compute the correlation between these two arrays of gene-gene correlation coefficient. (d) The gene expression levels of four common genes for the raw spatial data. (e) The gene expression levels of four common genes for the imputed spatial data.

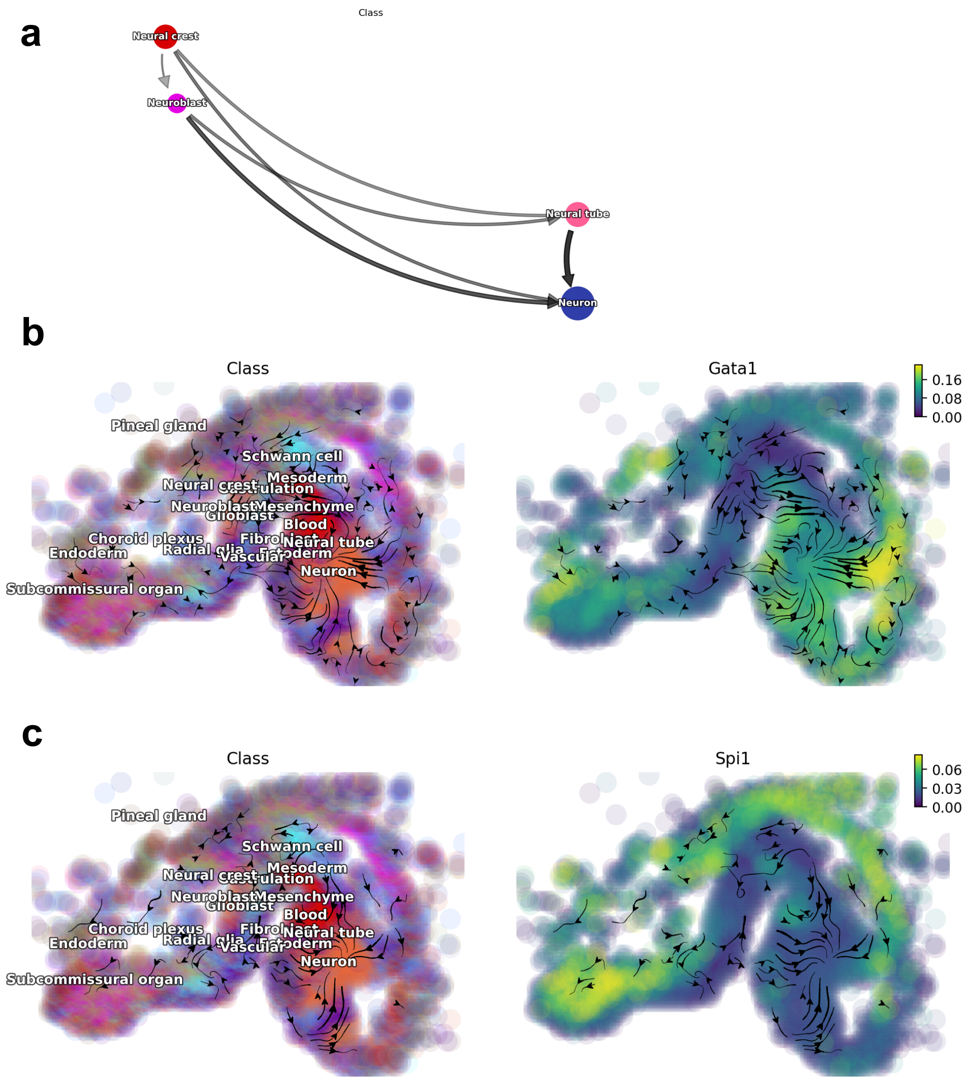


**Extended Data Fig. 12** In-silico perturbation for spatial data with dynamo. (a) The cell-type state transition graph of neural cells in the HybISS dataset after imputation. (b) RNA velocity after perturbing gene **Gata1**. The left panel represents the RNA velocity of spatial data colored by cell types. The right panel represents the RNA velocity of spatial data colored by the gene expression level of **Gata1**. (c) RNA velocity after perturbing gene **Spi1**. The left panel represents the RNA velocity of spatial data colored by cell types. The right panel represents the RNA velocity of spatial data colored by the gene expression level of **Spi1**.

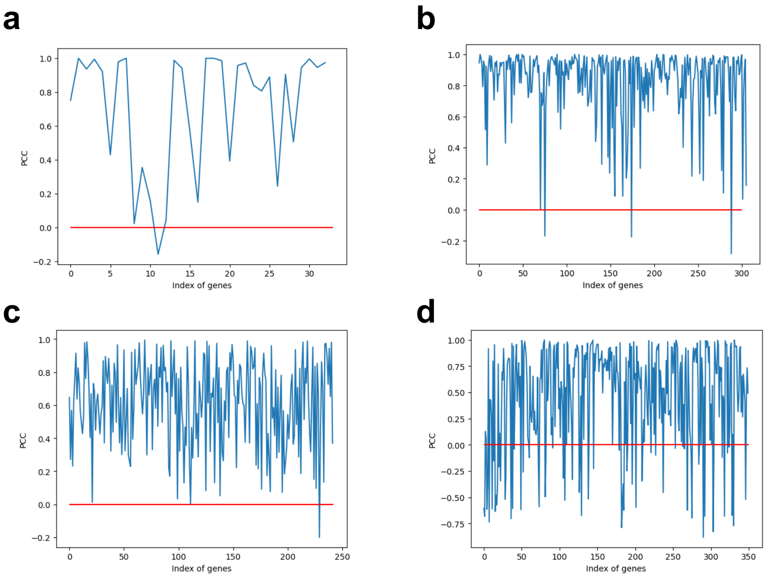


**Extended Data Fig. 13** PCC for the spatial data and scRNA-seq data. The y-axis represents the value of PCC and the x-axis represents the index of common genes. (a) PCC for the osmFISH-brain dataset and the reference scRNA-seq dataset. (b) PCC for the Xenium-breast dataset and the reference scRNA-seq dataset. (c) PCC for the Xenium-brain dataset and the reference scRNA-seq dataset. (d) PCC for the seqFISH-embryo dataset and the reference scRNA-seq dataset.

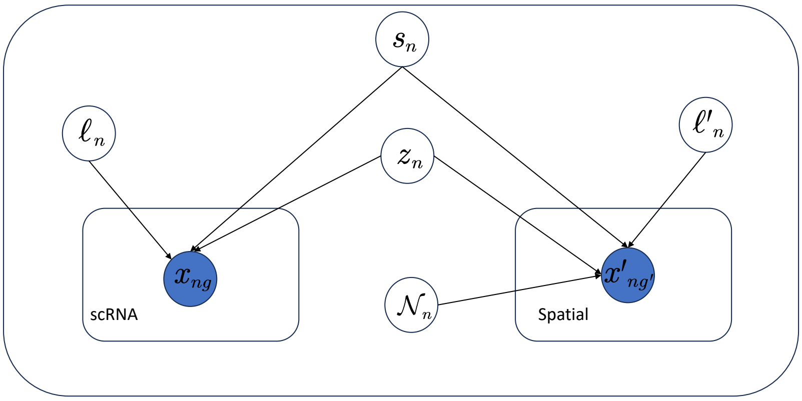


**Extended Data Fig. 14** Overview of our proposed mixture model. The definition of each symbol in this figure corresponds to the method section. scRNA-seq data and spatial data are formed into different domains with shared latent space containing biological information.

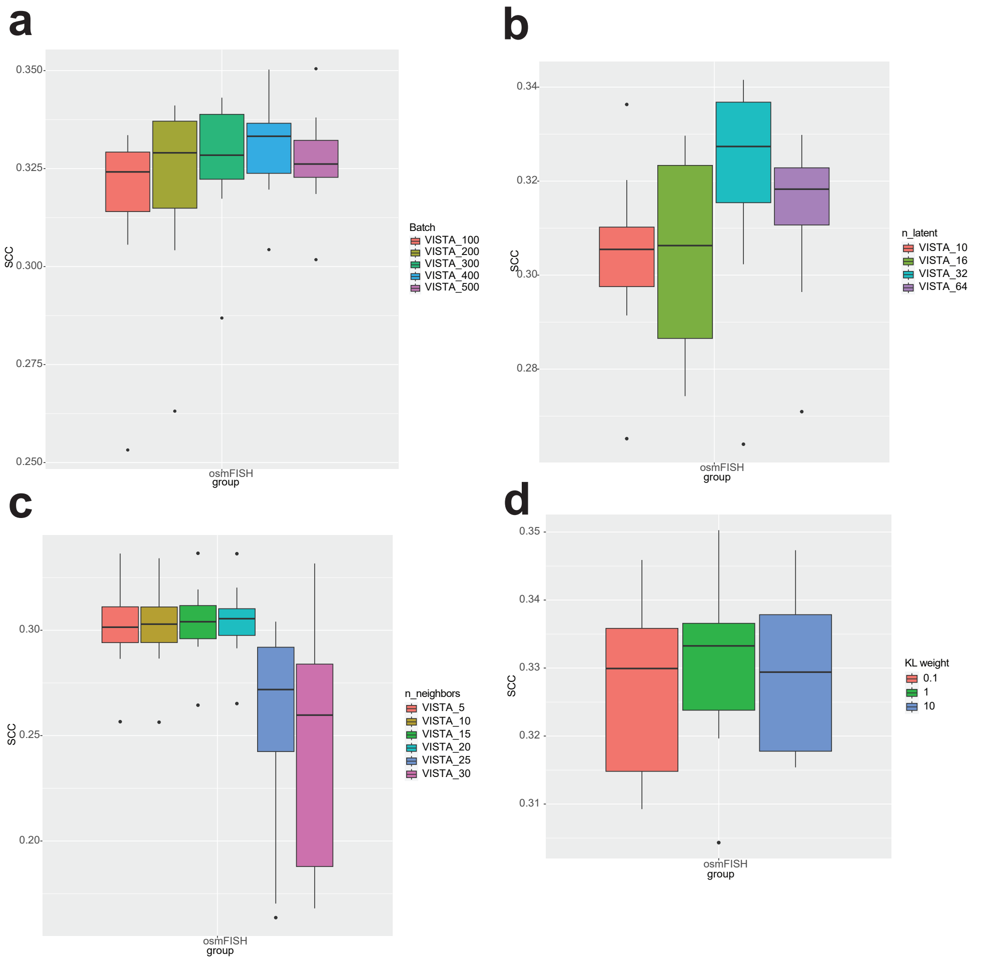


**Extended Data Fig. 15** Hyper-parameter tuning of VISTA based on osmFISH-brain dataset. (a) The change of SCC by adjusting epochs. (b) The change of SCC by adjusting the size of latent space. (c) The change of SCC by adjusting the number of neighbors. (d) The change of SCC by adjusting the weight for KL divergence in the loss function.

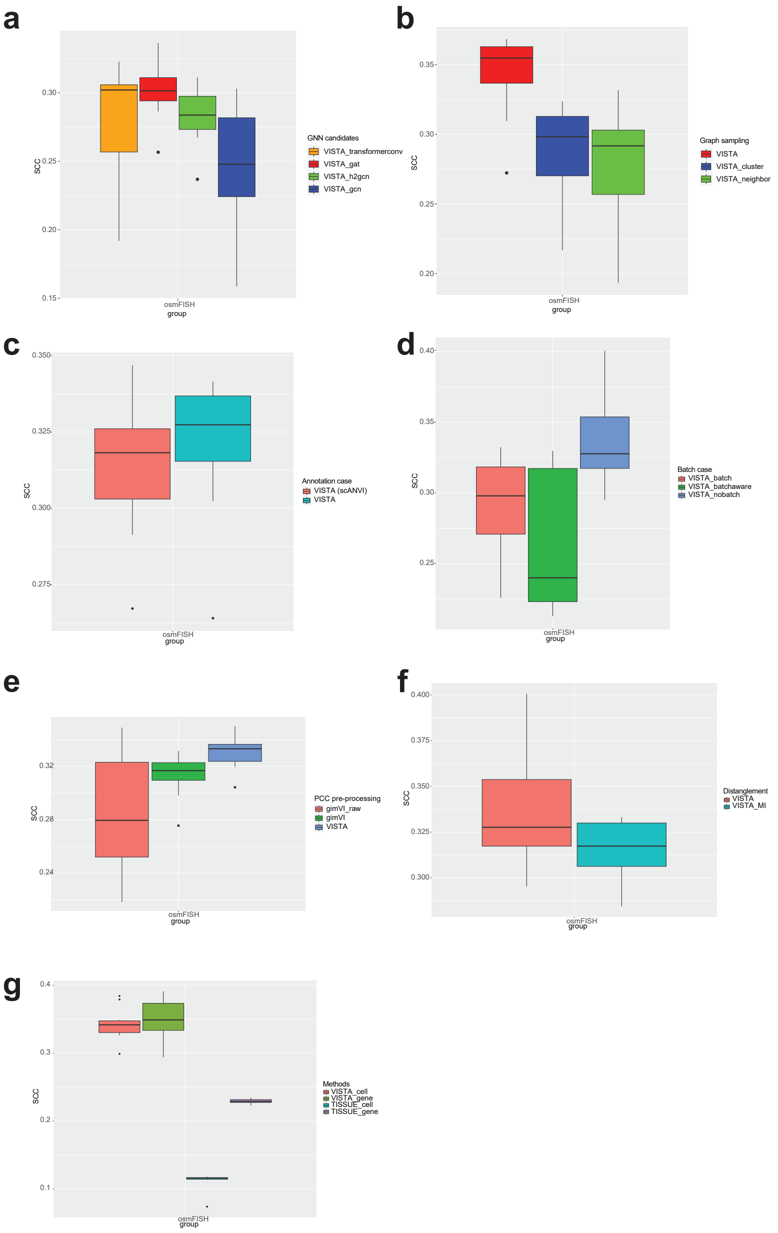


**Extended Data Fig. 16** Ablation test of VISTA based on osmFISH-brain dataset. (a) The change of SCC by adjusting the types of GNN we utilized. (b) The change of SCC by adjusting the choices of graph sampling methods. (c) The change of SCC by including cell-type information in the imputation or not. (d) The change of SCC by including batch-label information in the imputation or not. (e) The change of SCC by filtering genes with low correlation and significance levels or not. (f) The change of SCC by including mutual information in the loss function design or not. (g) The different results of SCC by changing the method for uncertainty estimation.

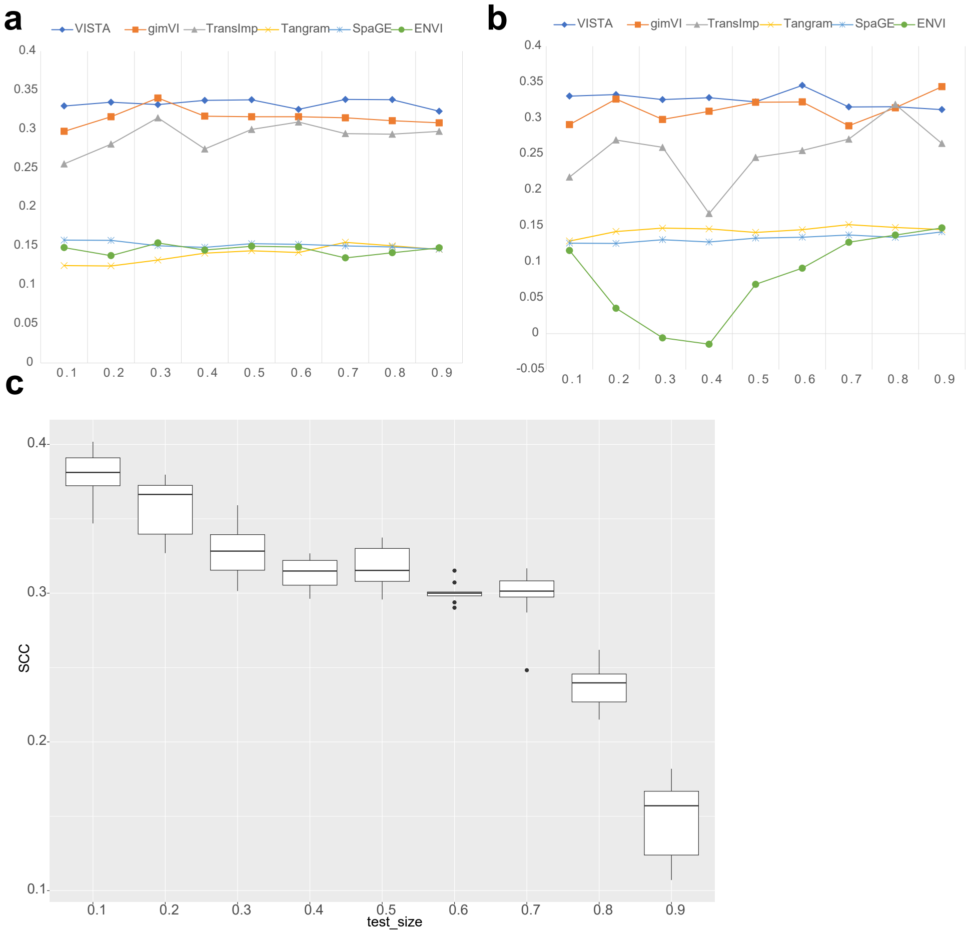


**Extended Data Fig. 17** Sensitivity analysis for the osmFISH-brain dataset. (a) SCC versus cell proportion of the spatial data across different methods. (b) SCC versus cell proportion of the reference scRNA-seq data across different methods. (c) SCC versus different testing sizes of genes.

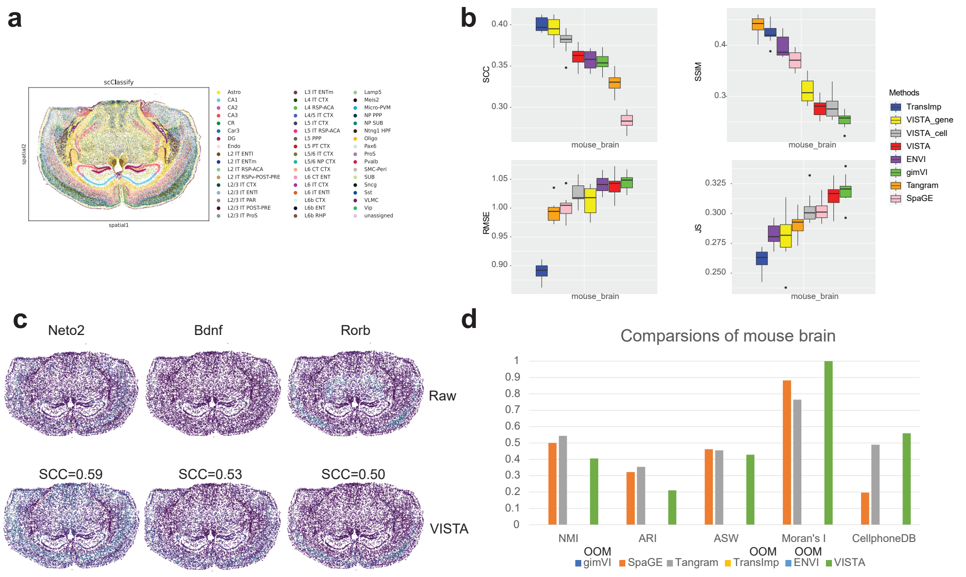


**Extended Data Fig. 18** Imputation results comparison for Xenium-brain dataset. (a) Cell-type distribution by spatial location of the Xenium-brain dataset. (b) Metrics information of different imputation methods. For SCC and SSMI, higher scores represent better performance, and the methods are sorted in descending. For RMSE and JS, lower scores represent better performance, and the methods are sorted in ascending. (c) Examples of reliable imputed genes produced by VISTA. The upper panel represents gene expression profiles in raw data, and the bottom panel represents gene expression profiles in the imputed data. (d) Metrics information of different imputation methods for Xenium-brain dataset.

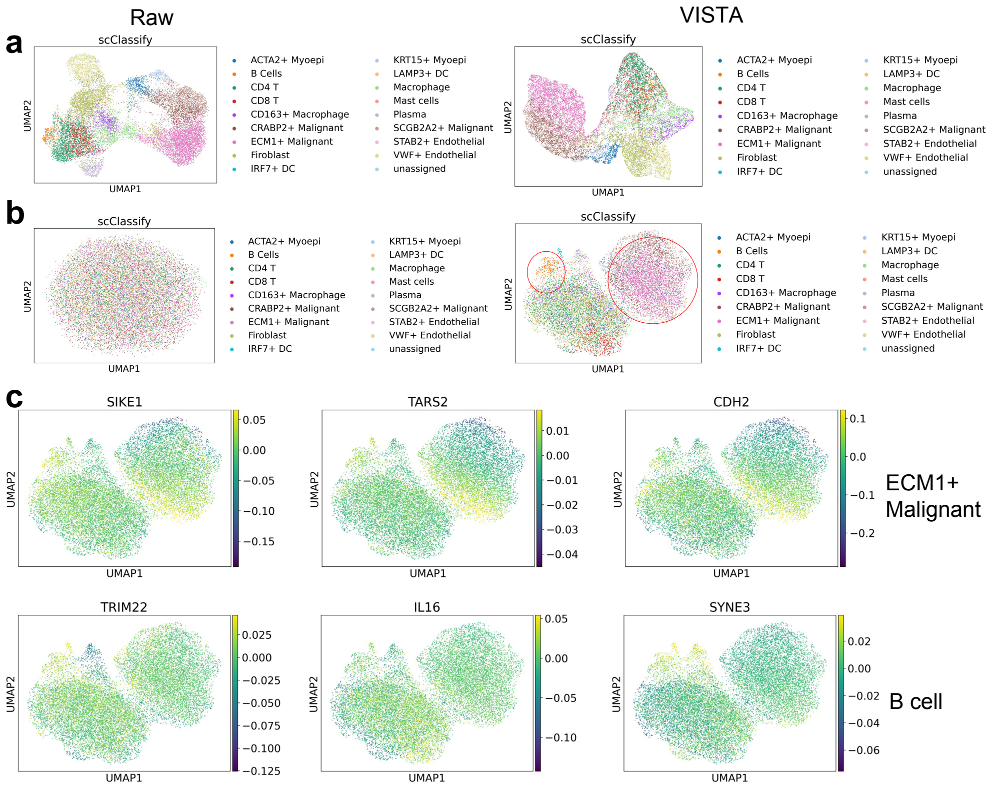


**Extended Data Fig. 19** Analysis of spatial variation of the Xenium-breast dataset. (a) UMAPs of the intrinsic variation colored by cell types for the raw spatial data and imputed spatial data. (b) UMAPs of the spatial variation colored by cell types for the raw spatial data and imputed spatial data. (c) The expression patterns of the DEGs are shown in the space of spatial variation. The upper figures represent DEGs of ECM1+ Malignant cells, and the bottom figures represent DEGs of B cells.
